# Simultaneous widefield voltage and interferometric dye-free optical mapping quantifies electromechanical waves in human iPSC-cardiomyocytes

**DOI:** 10.1101/2022.10.10.511562

**Authors:** Wei Liu, Julie Leann Han, Jakub Tomek, Gil Bub, Emilia Entcheva

**Affiliations:** Department of Biomedical Engineering, George Washington University, DC 20052, United States; Department of Computer Science, British Heart Foundation Centre of Research Excellence, University of Oxford, Oxford, United Kingdom and Department of Pharmacology, University of California, Davis, CA, United States; Department of Physiology and Physics, McGill University, Montreal, Canada; School of Electronic and Information Engineering, Harbin Institute of Technology, Shenzhen, China

## Abstract

Coupled electro-mechanical waves define heart’s function in health and disease. Genetic abnormalities, drug-triggered or acquired pathologies can disrupt and uncouple these waves with potentially lethal consequences. Optical mapping of electrical waves using fluorescent dyes or genetically-encoded sensors in human induced pluripotent stem cell derived cardiomyocytes (iPSC-CMs) offers mechanistic insights into cardiac conduction abnormalities. Interferometric dye-free/label-free wave mapping (without specific sensors) presents an alternative, likely capturing the mechanical aspects of cardiac conduction. Because of its non-invasive nature and spectral flexibility (not restricted to a specific excitation wavelength), it is an attractive chronic imaging tool in iPSC-CMs, as part of all-optical high-throughput platforms. In this study, we developed simultaneous widefield voltage and interferometric dye-free optical imaging methodology that was used: 1) to validate dye-free optical mapping for quantification of cardiac wave properties in human iPSC-CMs; 2) to demonstrate low-cost optical mapping of electromechanical waves in hiPSC-CMs using recent near-infrared (NIR) voltage sensors and orders of magnitude cheaper miniature CMOS cameras; 3) to uncover previously underexplored frequency- and space-varying parameters of cardiac electromechanical waves in hiPSC-CMs. We find similarity in the frequency-dependent responses of electrical (NIR fluorescence imaged) and mechanical (dye-free imaged) waves, with the latter being more sensitive to faster rates and showing steeper restitution and earlier appearance of wave-front tortuosity. During regular pacing, the dye-free imaged conduction velocity and the electrical wave velocity are correlated; both modalities being sensitive to pharmacological uncoupling and both dependent on gap-junctional protein (connexins) determinants of wave propagation. We uncover strong frequency-dependence of the electromechanical delay (EMD) locally and globally in hiPSC-CMs on a rigid substrate. The presented framework and results offer new means to track the functional responses of hiPSC-CM inexpensively and non-invasively for counteracting heart disease and aiding cardiotoxicity testing and drug development.

## INTRODUCTION

Electro-mechanical synchrony is critical for cardiac function. Under normal conditions, cardiomyocytes exhibit tight coupling between the electrical trigger (voltage), the calcium-induced calcium release and the mechanical contraction, known as excitation-contraction coupling, ECC(Bers 2002). In multicellular cardiac networks, electrical signals travel as fast synchronized waves. Due to the tight ECC, these voltage waves are closely followed by calcium waves; in fact, calcium waves are often used as a surrogate for the cardiac excitation due to better optical sensors for calcium. At the level of each cell, a mechanical contraction ensues following the electrical trigger and the beginning of the calcium transient. Therefore, mechanical waves, driven by the local EC coupling, spread over space similarly to the excitation waves. Macroscopically, the dynamics of this tri-wave system (voltage, calcium, contraction) is dictated by the fast-diffusing voltage waves. However, locally, the much slower diffusing calcium can exhibit uncoupled behavior with no correspondence to the global activation, e.g. microscopic local calcium waves can be confined to individual cells and can co-exist with the global activation(Aistrup, Kelly et al. 2006). The calcium-voltage dynamics on a global (wave) scale can be quite complex due to inherent feedback loops between calcium and voltage(Shiferaw, Sato et al. 2005, Shiferaw and Karma 2006), and this interplay can influence the risk for arrhythmia development. Similarly, the spread of mechanical activity as a wave of contraction can show significant deviations from the excitation wave – it can precede it (mechano-electrical event), or it can deviate in speed and in morphology from the excitation event due to mechanical loading conditions and the level of mechanical coupling between the cells, which involve non-local elastic interactions(Nitsan, Drori et al. 2016, Pfeiffer-Kaushik, Smith et al. 2019). Because of feedback mechanisms (e.g. mechano-sensitive ion channels), significant deviations in the patterns of mechanical contraction waves can, in turn, further influence the stability of the excitation waves. The overall dynamics of the tri-wave system can be very complex and hard to map out, yet these interactions are of great importance in understanding, treating and preventing cardiac arrhythmias.

Cardiac cell culture systems represent a reductionist approach to dissect wave dynamics properties. Early macroscopic optical mapping using calcium and voltage sensitive fluorescent dyes in such two-dimensional systems confirmed the existence of reentrant waves in culture, with fundamental properties resembling those seen in the whole heart(Bub, Glass et al. 1998, Entcheva, Lu et al. 2000, Arutunyan, Webster et al. 2001, Iravanian, Nabutovsky et al. 2003). With the emergence of human stem-cell-derived cardiomyocytes, especially induced pluripotent stem-cell-derived cardiomyocytes, iPSC-CMs, such models became of interest(Weinberg, Lipke et al. 2010, Herron, Lee et al. 2012, Gintant, Burridge et al. 2019) as pre-clinical tools in drug development and as means towards personalized medicine.

In addition to optical mapping of excitation, optical contraction measurements by variants of frame differencing have been used under different names, reflecting conceptually similar approaches: video tracking, digital cross-correlation, digital image correlation, or “optical flow” on the surface of a whole heart(Gaudette, Todaro et al. 2001, Zhang, Iijima et al. 2016, Christoph and Luther 2018) or in monolayers of cardiomyocytes (Peeters, Hlady et al. 1987, Bien, Yin et al. 2003, Entcheva and Bien 2003, Chung, Bien et al. 2011), including applications to human iPSC-CMs (Ahola, Kiviaho et al. 2014, Huebsch, Loskill et al. 2015, Pointon, Harmer et al. 2015, Hansen, Favreau et al. 2017, Sala, Meer et al. 2018). These methods, applied at the microscale (small field of view) have been used to track uniaxial strains(Peeters, Hlady et al. 1987, Bien, Yin et al. 2003) or biaxial strain fields(Gaudette, Todaro et al. 2001, Chung, Bien et al. 2011). In some variants, spectral domain methods have been applied to track periodic endogenous structures, such as sarcomeres, or to track the deformation of imposed light interference patterns (Moire fringes)(Andonian 1984, Zheng and Zhang 2008, Zheng, Surks et al. 2010). Such systems, developed to analyze the functional responses of human iPSC-CMs in drug testing, have the ability for sequential or simultaneous triple measurements (voltage, calcium and contraction), sometimes also combined with optogenetic pacing (Hortigon-Vinagre, Zamora et al. 2016, Klimas, Ambrosi et al. 2016, van Meer, Krotenberg et al. 2019, Klimas, Ortiz et al. 2020).

However, dye-free/label-free macroscopic optical mapping of contraction waves (large field of view) has been demonstrated only in a few studies in cultured cells. For example, Hwang et al. (Hwang, Yea et al. 2004, Hwang, Kim et al. 2005) used on-axis white light trans-illumination with a pinhole to demonstrate the first dye-free tracking of complex cardiac contraction waves in cultured cardiomyocytes. In a similar setup, termed “propagation-induced phase contrast imaging”, Rother et al. (Rother, Richter et al. 2015) quantified contraction waves in cardiomyocytes-fibroblasts co-cultures. Burton et al. used oblique trans-illumination with a semi-coherent light source for dye-free imaging of cardiac waves while applying optogenetic control patterns(Burton, Klimas et al. 2015) and to study neuro-cardiac effects on wave dynamics(Burton, Tomek et al. 2020).

A versatile yet simple methodology for simultaneous tracking of electrical and mechanical waves in cardiac cells and tissues is of great interest. Developing such a system for human iPSC-cardiomyocytes is the objective of the current study. In a simplified setting, it can provide fundamental understanding about the interaction of these co-existing dynamical systems and the conditions under which their interaction may be destabilizing and pro-arrhythmic. Obtaining comprehensive multiparametric assessment in human iPSC-CMs can lead to better experimental pre-clinical models for drug development and personalized medicine(Gintant, Burridge et al. 2019, Yang, Ribeiro et al. 2022). Such knowledge can also improve *in silico* electromechanical models, which have been developed over the years(Nash and Panfilov 2004, Trayanova, Constantino et al. 2011, Trayanova and Rice 2011, Wang, Santiago et al. 2021) but not extensively validated. Dissecting such wave interactions can inform translational imaging of electromechanical waves in the heart for noninvasive arrhythmia mapping in the clinic(Provost, Lee et al. 2011, Christoph, Chebbok et al. 2018, Grubb, Melki et al. 2020).

## RESULTS

In this study we set out to develop means for the simultaneous tracking of electrical and mechanical waves in a simplified system of heart tissue – an isotropic syncytium of human iPSC-cardiomyocytes, attached to a hard substrate. A reduced experimental cardiac model offers a well-controlled starting point to quantify the properties of these co-existing waves before adding aspects of the complexities of mechanical loading in the real heart.

### Simultaneous dual macroscopic imaging (SDMI) system for electromechanical waves

The SDMI system presented here (**Figure 1A**) uses oblique transillumination to simultaneously map voltage (by fluorescence) and mechanical waves (by dye-free signal) in human iPSC-CMs at the macroscale (large field of view, FOV). The oblique-light interferometric tracking of waves without dyes is based on a technique described in our previous work(Burton, Klimas et al. 2015); it is believed that the captured signal is related to local changes in the optical path length as the cells contract. Because of the attachment to fibronectin-coated glass, the cell movements are localized micro-contractions. In this study, two low-cost (< $500) miniature CMOS cameras were used to simultaneously track the electrical and mechanical waves in cardiac monolayers (**Supplemental Video 1**) by fast optical mapping at high spatial resolution. Here, we show that they are sensitive enough to yield comparable performance to state-of-the-art scientific sCMOS, e.g. Photometrics 95B, in optical recordings of voltage and calcium (by fluorescence) and of contraction (dye-free) in human iPSC-CMs at the microscale (**Figure S1, S2**) and at the macroscale. Synchronized acquisition of the two cameras was controlled from the same trigger sequence by a function generator. Light sources with relatively long temporal coherence length (∼20 μm; comparable to cell size) were used. In our implementation this was either a laser diode or a red LED. Oblique transillumination helped capture the scattering interference patterns (by the dye-free camera) with minimal incident beam background (similar to dark-field imaging), which has little effect on the fluorescence-based voltage mapping. Additionally, oblique transillumination yields higher spatial resolution over direct illumination for interferometric imaging (Junger, Olshausen et al. 2016).

**Figure 1.**
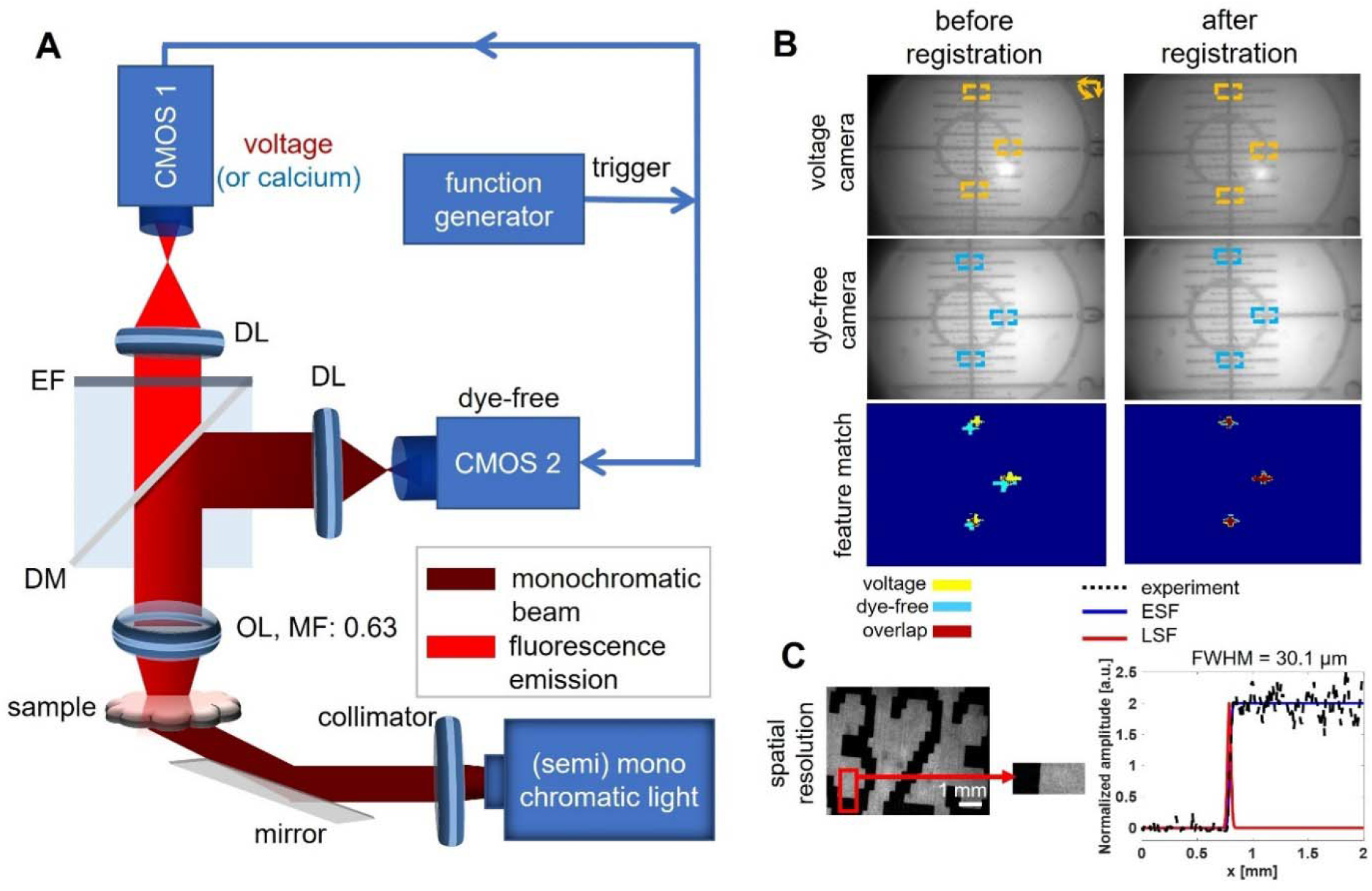
Simultaneous widefield (NIR fluorescence) voltage and (interferometric) dye-free optical mapping of electromechanical waves in hiPSC-CMs using oblique transillumination and low-cost CMOS cameras: experimental setup, camera co-registration and spatial resolution. A. Schematic diagram of the macroscopic imaging system with two Basler CMOS cameras synchronized by a trigger. A laser or an LED (660nm) were used for oblique illumination at 30 degrees. At 0.315x magnification, the FOV was about 16×12 mm^2^. B. Dual-macroscopic images of a pattern before and after co-registration by feature-extracted match (see also **Figure S2** and **Supplement**). The extracted cross features were labeled by the dashed boxes. C. Lateral spatial resolution measured by the full width at half max (FWHM) of the line spread function of high-resolution printed letters was 30.1μ m. Measured area was marked by the red rectangle. DM: dichroic mirror; DL: demagnification lens; EF: emission filter; ESF: edge spread function; LSF: line spread function; MF: magnification factor; OL: objective lens.

The electrical waves were mapped with a high-performance NIR voltage-sensitive dye Berst1(Huang, Walker et al. 2015), which we have used before for optical mapping in iPSC-CMs(Klimas, Ortiz et al. 2020). The emitted fluorescence and the dye-free signal were imaged by a low-NA objective (NA: 0.15) and split to the corresponding cameras by a dichroic mirror. With an additional demagnification, the voltage and dye-free cameras shared the FOV with a slight difference, **Figure 1B**. Co-registration of the two cameras was done by feature-extracted match of a pattern (see **Figure 1, Figure S3** and **Supplemental Methods**) to guarantee precise comparison of the two imaging modalities. The spatial resolution of the two imaging modalities was quantified as 30.1 μm by measuring the full width at half max (FWHM)(Liu, Shcherbakova et al. 2018) of the line spread function of high-resolution printed letters (**Figure 1C**). The two imaging modalities hold the same imaging resolution due to shared optical path.

### Initiation of electromechanical waves in human iPSC-CMs, data processing, validation of the dye-free mapping and extensibility

Cardiac arrhythmias manifest as electromechanical waves that are self-driven and follow complex paths and dynamics. To reduce complexity and to uncover general guiding principles for cardiac conduction, experimental models of the heart, including human iPSC-CMs, often resort to studies that explore the restitution properties of the system, i.e. responses to different pacing rates in controlled conditions. Similarly, here, we analyzed spontaneous and controlled paced activity, to begin to understand the basics of electro-mechanical wave dynamics. Pacing was done electrically using a bipolar point electrode; the response to a range of pacing frequencies was quantified for each sample in addition to spontaneous waves. In our microscopic scale validation experiments, see **Figures S1, S2**, as well as in a companion manuscript(Heinson, Han et al. 2022), we used optogenetic pacing, coupled with optical mapping of voltage, calcium and/or contraction for all-optical interrogation of cardiac electromechanics, inspired by previous studies (Arrenberg, Stainier et al. 2010, Burton, Klimas et al. 2015, Klimas, Ambrosi et al. 2016, Klimas, Ortiz et al. 2020, Heinson, Han et al. 2022).

The mapping data for the electrical and mechanical waves, captured by the two cameras, were processed using empirically optimized methods to extract a multitude of parameters and metrics, **Figure S4**. Signal to noise ratio (SNR) was improved by taking advantage of the periodic nature of mapped activity in this study and using a period-shift-enhancement (PSE) as described in the Methods. Because of the slower upstroke rate of cardiac motion (in contrast to the sharper depolarization in the action potential), the automatic detection of the motion start point presents a challenge. Our solution to overcome the noise in the dye-free signal and to boost event detection was to use a time-difference (TD) filter (**Figure S4-S5**), as done previously (Burton, Klimas et al. 2015). The signal enhancement of the dye-free traces by the TD filter can be appreciated in the microscopic records in **Figure S2**. For quantification of local parameters and relative characteristics of the electro-mechanical waves, instead of the TD boost, we used the original dye-free signal and an alternative SNR improvement-activation-based spatial segmentation, see a branch in the signal processing pipeline in **Figure S4**. This helped obtain reliable action potential duration, APD, dye-free signal duration (DFD), relaxation-contraction ratio, and, importantly, apparent electro-mechanical delay, aEMD. After spatial and temporal filtering and event detection (determination of local activation time), conduction could be visualized by activation maps and quantified by conduction velocity (CV) measurements and wavefront tortuosity (WT) index measurements, **Figure S4**.

To validate the dye-free mapping of cardiac waves in our system, we first used label-free samples of hiPSC-CMs to record the optical signal under different pacing conditions, as shown in **Figure 2A**. Expected wave origin at the pacing site and slowing of conduction velocity with increased pacing frequency are illustrated using activation maps derived from the dye-free signal. Furthermore, even though the rest of this study was conducted using dual voltage – dye-free mapping in controlled paced conditions, in **Figure 2B** we demonstrated that the dye-free imaging is extendable to calcium – contraction dual optical mapping. This was done using a model of anatomical reentry – a non-paced self-sustained activation around a non-excitable obstacle/annulus (in this case, removed portion of the cell layer). See the Methods and companion manuscript(Heinson, Han et al. 2022) for details on the changes needed to accommodate calcium imaging.

**Figure 2.**
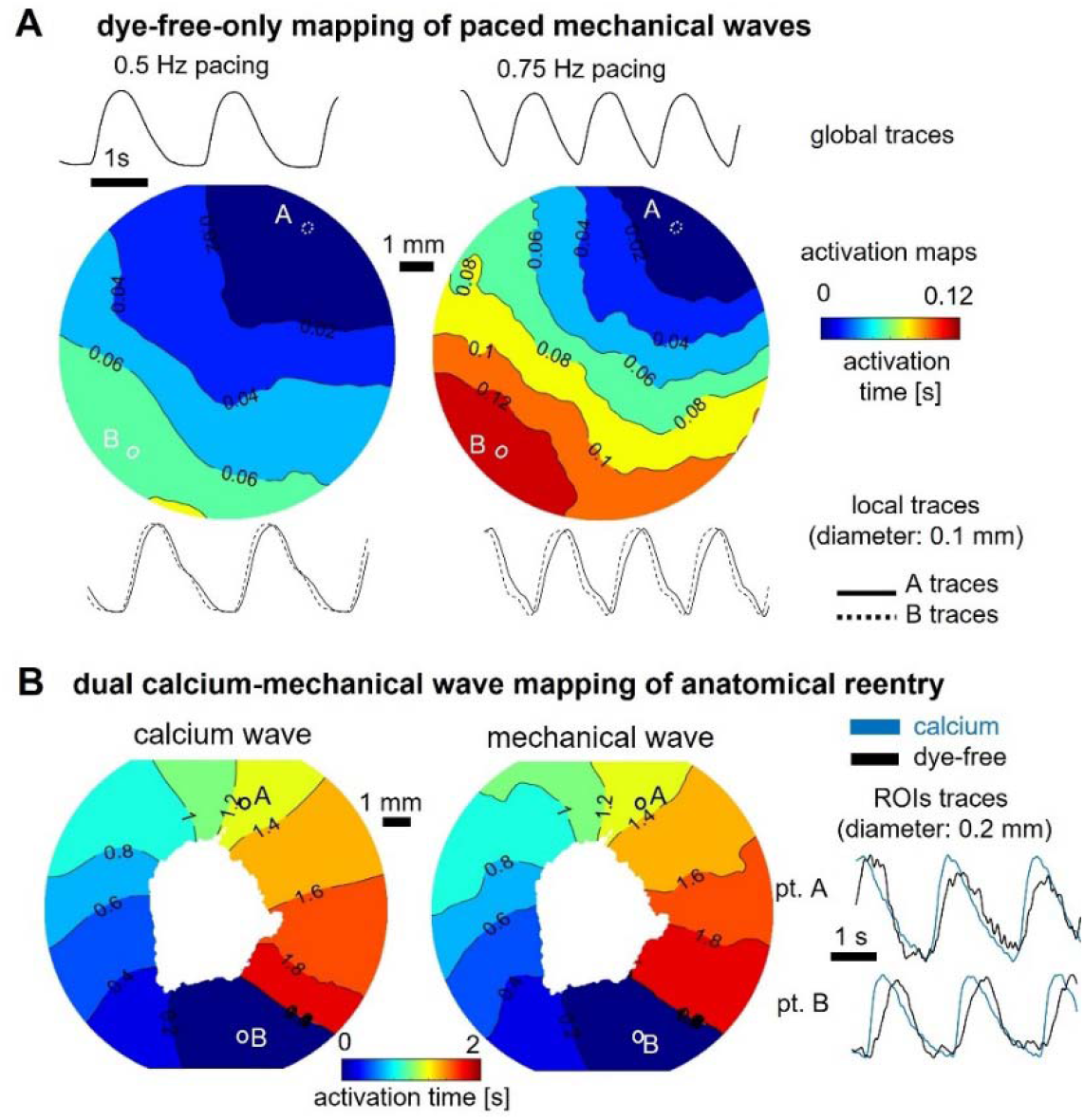
Dye-free only optical mapping and simultaneous calcium - dye-free widefield imaging of reentrant waves. A. Optical mapping of paced mechanical waves in unlabeled hiPSC-CMs using oblique illumination and interferometric dye-free imaging: activation maps, global and local traces shown. B. Simultaneous optical mapping of calcium waves and mechanical waves using the system in Figure 1 with appropriate light sources and filters; samples labeled with Rhod-4 calcium-sensitive dye. Anatomical reentrant wave was induced by removing cells from the center of the dish to create an obstacle and applying rapid pacing until a sustained self-propagating reentrant wave was observed.

### Similarities and differences of electrical and mechanical waves in human iPSC-CMs

Key parameters were quantified for the electrical and mechanical waves globally and locally, under different pacing conditions. The global traces, over a large FOV, **Figure 3A**, are akin to clinical ECG or global ultrasound-detectable contraction waves and help confirm the 1:1 response to pacing. Local traces help map activation patterns in detail and visualize the waves. **Figure 3B** illustrates that both electrical and mechanical waves exhibit CV restitution, i.e. wave slowing with increase in frequency. The mechanical waves in human iPSC-CMs are substantially more sensitive to progressively increasing pacing rates, with abnormalities in conduction and slower regions appearing earlier than in the voltage-based maps. Furthermore, the spatial patterns of wavebreaks appearing first in the mechanical maps may be predictive of the location of electrical wavebreaks at a higher frequency point, as seen in **Figure 3B**. These are important observations for early detection of arrhythmias clinically because three-dimensional high-resolution mechanical mapping is easier to obtain non-invasively compared to obtaining unambiguous high-resolution electrical mapping (Grondin, Costet et al. 2016, Christoph, Chebbok et al. 2018). Dye-free imaging may also be a suitable highly sensitive tool for arrhythmia detection in human iPSC-CMs in scalable cardiotoxicity testing platforms(Burton, Klimas et al. 2015, Entcheva and Kay 2021).

**Figure 3.**
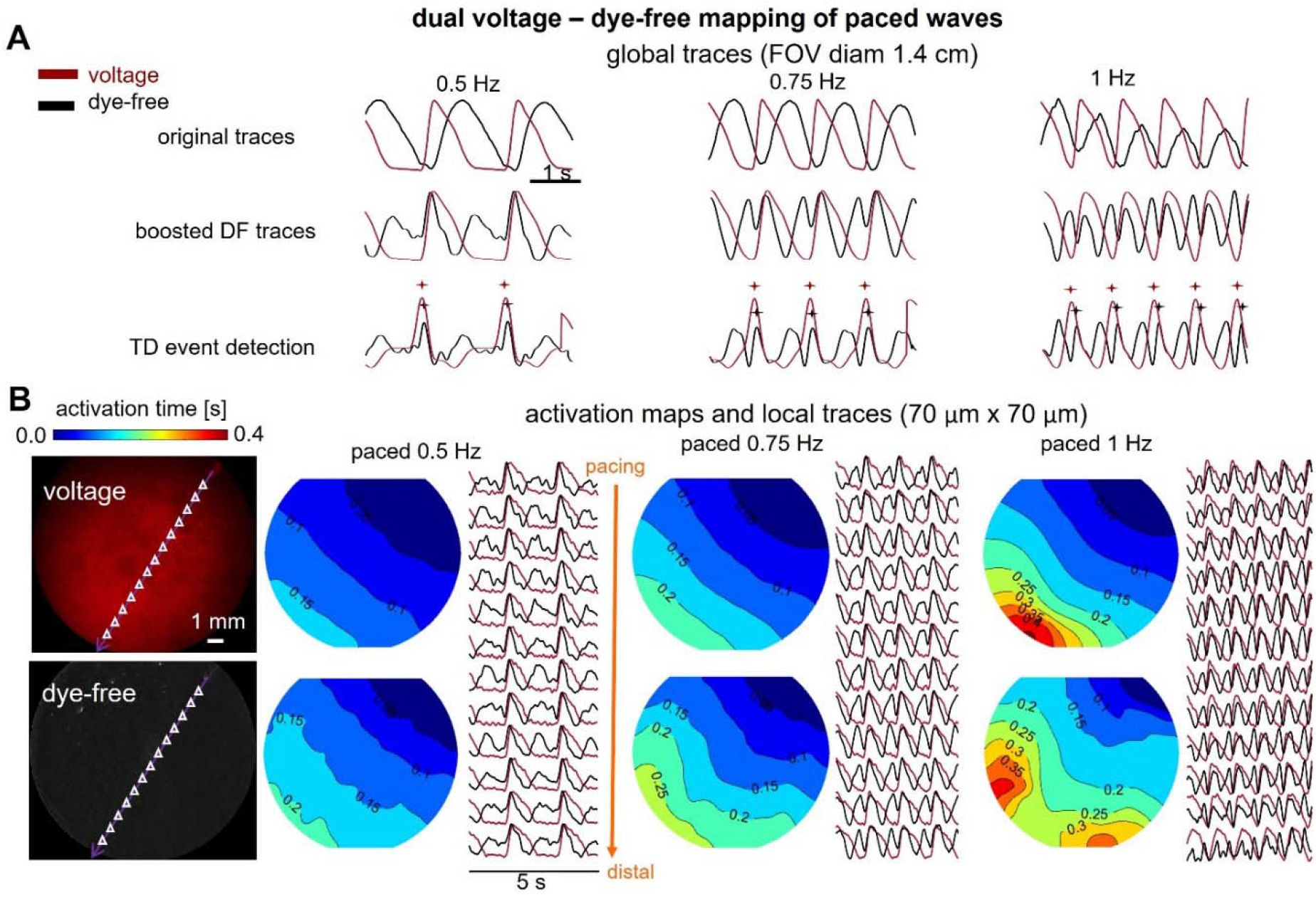
Comparison of activation maps from simultaneous widefield-imaged voltage and dye-free signals in hiPSC-CMs under different stimulation frequencies. A. Normalized global (averaged over the FOV) action potentials imaged by the NIR voltage-sensitive dye BeRST1 (red) and dye-free signals - original and TD-boosted (black) under 0.5 Hz, 0.75 Hz and 1 Hz pacing. Shown at the bottom are the respective detected activation times (marked) by TD filter. B. Macroscopic NIR fluorescence (top) and dye-free (bottom) interferometric images of the hiPSC-CMs samples in the same FOV (left). Small triangles indicate the centers of the matched regions of interest (ROIs, 70 μm × 70 μm each) for the local trace plots. Shown on the right are voltage (top) and dye-free (bottom) activation maps along with the extracted local traces at the indicated triangle points under 0.5 Hz, 0.75Hz and 1Hz pacing from a point in the upper right corner. Isochrones are 0.05s apart. All measurements were done at room temperature.

A detailed quantitative analysis of electromechanical wave properties (**Figure 4**) offers further insights related to wave dynamics and emergence of instabilities. The mechanical DFD exhibits a much wider range of durations and steeper restitution compared to the APD, **Figure 4A**. This rate adaptation in the mechanical signal comes primarily from change in the relaxation phase, not the contraction phase, **Figure 4B**. Overall, APD/DFD ratios steadily increase with frequency, approaching a ratio of 1, **Figure 4C**. From our results, it appears that DFD changes, obtainable via dye-free optical mapping, may be useful as a metric to capture restitution responses locally. As suggested in **Figure 3**, and quantified in **Figure 4D-F**, both electrical and mechanical waves exhibit CV restitution, yet they dramatically differ in the effects of rate on wavefront morphology and stability. We introduce a wavefront tortuosity index, WT index, to quantify the significantly higher proneness of mechanical waves to wavebreaks and local instabilities, **Figure 4E-F**, compared to voltage-based conduction. The higher tortuosity of the mechanical wave front reflects variation in cellular excitation-contraction coupling or variation in the mechanical resistances of adjacent cells after contraction is initiated (mechanical coupling) or both. The effect of any coupling heterogeneity will naturally slow down the mechanical wave and will disturb the wavefront(Gokhale, Asfour et al. 2018). To some degree, the difference in the WT index reveals the difference in the local electrical coupling efficiency for voltage wave propagation vs. the local mechanical coupling efficiency (associated with cellular excitation-contraction coupling and mechanical resistance) for mechanical wave propagation. A larger WT index for the mechanical waves represents lower coupling efficiency, further exacerbated by stressors like faster pacing rates. The wavefront stability and the local CV can be vastly different between the two mapping modalities, **Figure 4F** and **Figure S6**. Overall, despite similarities in the global wave properties between electrical and mechanical waves, the mechanical wave dynamics exhibits higher local complexity and level of instabilities compared to electrical waves over space and time.

**Figure 4.**
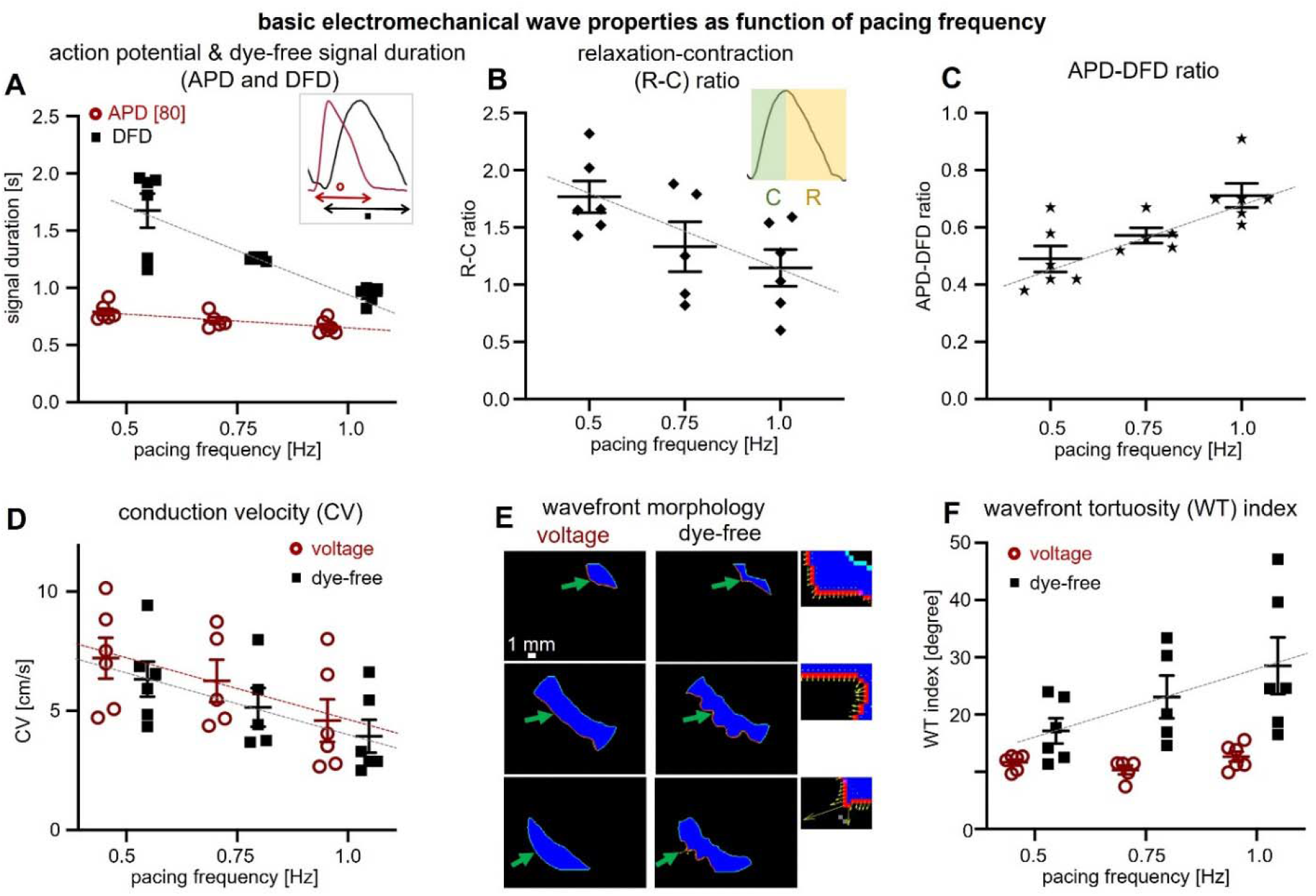
Quantification of basic parameters of electromechanical waves in hiPSC-CMs by simultaneous widefield voltage and dye-free optical mapping. A. Restitution (frequency response) of action potential and dye-free signal duration (APD & DFD) for all samples; voltage in red (slope: −0.26; R^2^: 0.45) and dye-free in black (slope: −1.48; R^2^: 0.7). Inset shows that for voltage the measurement was at 80% level, i.e. APD80, and for the original dye-free signals the transient duration measurement was between the start of contraction and the end of the relaxation. B. Frequency response of the relaxation-contraction (R-C) ratio (see inset) for all samples based on dye-free signals (slope: −1.24; R^2^: 0.33). C. Electromechanical signal duration ratio, i.e. APD-DFD ratio, as a function of pacing frequency (slope: 0.44; R^2^: 0.52). D. Conduction velocity (CV) restitution (voltage & dye-free slopes: −5.24 & −4.78 and R^2^: 0.25 & 0.28, respectively. E. Wave-front morphology-voltage and dye-free wave-front segmentation from an activation map (obtained at 0.5-Hz pacing) using frames that are 0.15s apart. Close-up insets showing the local dye-free velocity vectors; green arrows indicate the wavefront. F. Wave-front tortuosity (WT) index (for details see Methods) quantified for voltage and dye-free waves for all samples as a function of the pacing frequency (dye-free slope: 22.77; R^2^: 0.25). All measurements were at room temperature. Data are presented as scatter plots (mean values per sample) with overlaid mean ± S.E., n = 6 samples; all shown overlayed regression lines (dependence on frequency) passed the significance test at p<0.05; the WT index for voltage did not have a significant dependence on frequency.

### Correlation of electrical and mechanical wave properties to gap junctional protein, Cx43 and response to pharmacological uncoupling

Because of strong excitation-contraction coupling in cardiomyocytes, the global CV for the electrical waves is closely correlated with the global CV for the mechanical waves, see **Figure 5A**, summarizing the data from **Figure 4D**. Thus, global CV obtained from dye-free wave mapping under regular periodic activity can likely be used as a surrogate of the global CV of the electrical wave. This does not hold true for the local CV – mechanical waves exhibit much higher local heterogeneity in conduction compared to electrical waves (**Figure 4E-F**). The transient durations - DFD and APD - exhibit lower degree of correlation to each other compared to the electromechanical wave CVs (**Figure 5B**). DFD is much more sensitive to restitution and the strongest frequency-dependent adaptation happens via shortening of the mechanical relaxation phase and the DFD. The least correlated parameter between the two waves is the WT index in response to pacing frequency – while the mechanical wavefront is acutely influenced by the pacing stressor, the electrical wavefront remains stable (**Figure 5C**). The key determinant of the global electrical wave conduction velocity is the electrical coupling through dedicated gap junctional channels. In ventricular cardiomyocytes, including human iPSC-CMs, the dominant gap junctional protein is Connexin 43 (Cx43). We sought to pharmacologically inhibit gap junctional coupling with heptanol (0.5mM for 15min) and quantify its effect on both waves. In the example in **Figure 5D-E**, the gap junctional inhibition resulted in about 17% reduction in CV for both waves, with some non-specific shortening in APD and DFD, which was more pronounced for APD, leading to reduction in APD-DFD ratio. The most dramatic effects of heptanol were on the aEMD. The average local aEMD was increased by about 30% and the increase in the global aEMD was close to 50%. Therefore, gap junctional defects or exogenous uncouplers may lead to arrhythmias not only through pro-arrhythmic slowing of electrical wave CV but also through uncoupling of the electrical and mechanical waves.

**Figure 5.**
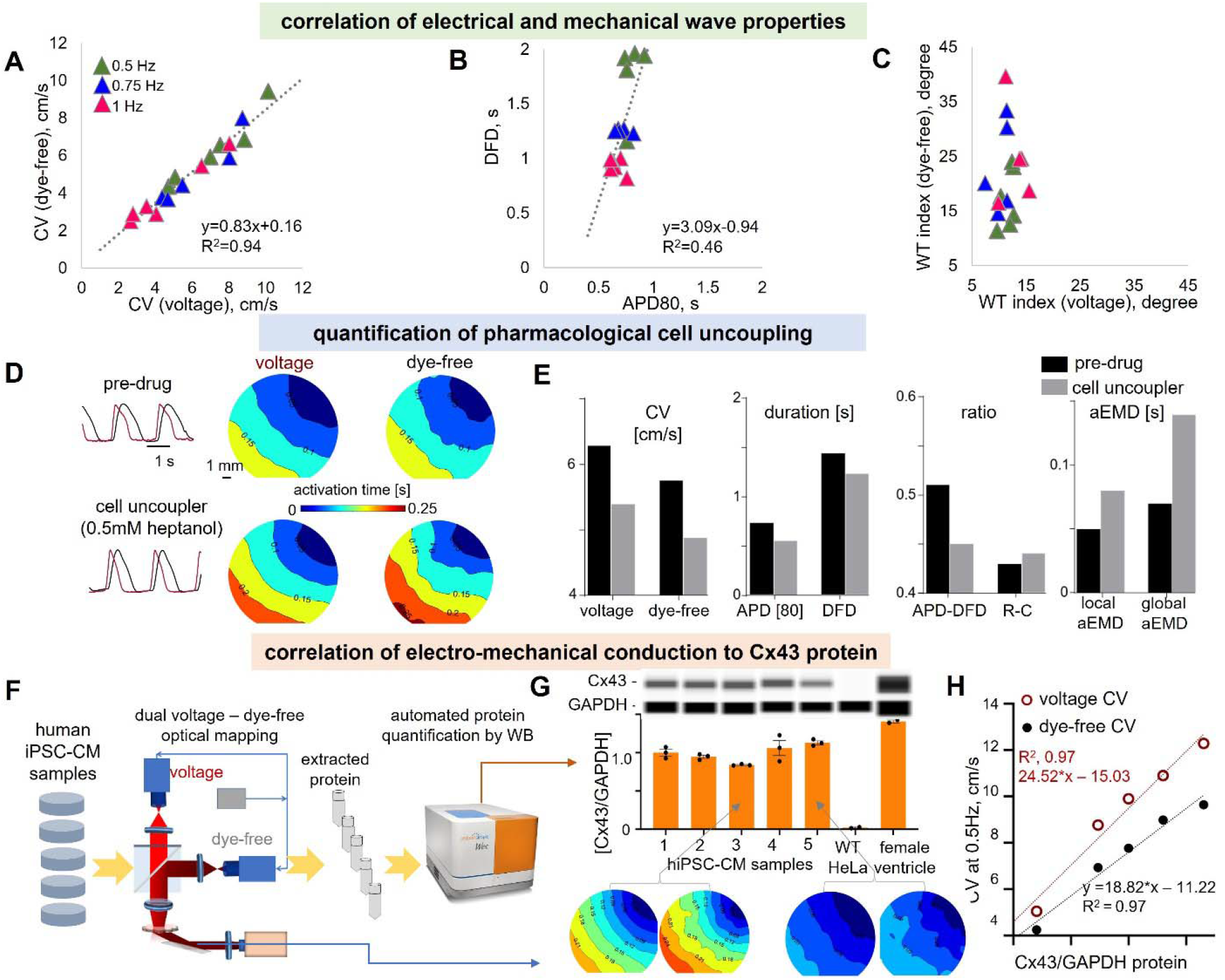
Correlation of electrical and mechanical conduction properties, ability of voltage and dye-free imaging to detect pharmacologically-inhibited conduction and correlation of electromechanical conduction to gap junctional (Cx43) protein levels. A-C. Correlation of electromechanical properties for individual samples, n = 6: A. Voltage – dye-free correlation of CV across pacing frequencies. Linear regression was applied for data fitting (slope: 0.83; R^2^: 0.94). B. Voltage – dye-free correlation of transient duration DFD vs APD across pacing frequencies, linear regression line (slope: 3.09; R^2^: 0.46). C. Voltage – dye-free correlation of WT index across pacing frequencies. D-E. Pharmacological inhibition of conduction by cell uncoupler heptanol and the response seen by voltage and dye-free optical mapping, n=1. In panel D, shown are activation maps from voltage and dye-free optical mapping pre-drug and after 15 min of 0.5mM heptanol; shown on the left are global voltage and dye-free signals. In panel E, various parameters from the dual imaging are compared pre-drug and after heptanol application for the voltage signals (CV, APD) and for the dye-free signals (CV, DFD). Furthermore, the heptanol-triggered change in electro-mechanical coupling parameters was quantified, e.g. APD-DFD ratio, R-C ratio and average local aEMD and global aEMD. Both modalities sense cell-uncoupling in a similar way; electromechanical delay is increased by cell-uncoupling, most pronounced as a prolongation of the global aEMD. F-H. Correlation of electromechanical conduction to Cx43 protein levels, n=5 (two of the samples are from the same set as in Figure 4, and three of them are additional samples). Panel F presents a pipeline for correlating functional properties to Cx43 protein by Western Blot (WB). After functional optical measurements, protein was extracted, and WB was done by the automated capillary electrophoresis Wes system, keeping track of the sample identity. Panel G shows the Wes-generated digital lanes (obtained from electropherograms) for antibody-multiplexed Cx43 and a housekeeping protein GAPDH, along with the quantified ratios for the five samples from several runs / technical replicates. Negative and positive controls for Cx43 were lysates from WT HeLa cells and from human female heart ventricular tissue. Underneath, in panel G, are shown activation maps for voltage and dye free for sample 3 and sample 5. Panel H shows individual sample correlation of CV for voltage (red) and dye-free (black) to the Cx43/GAPDH ratios. Linear regression lines for voltage showed slope of 24.52 and R^2^ = 0.97, and for the dye-free signal slope of 18.82 and R^2^ = 0.97. The Wes image was generated using Biorender software.

We developed a pipeline to link functional wave properties and gap junctional protein quantity in the same samples, as shown in **Figure 5F**. Five samples were imaged optically using the SDMI system and then cells were lysed and protein extracted and quantified using capillary-based electrophoresis (Wes™ by ProteinSimple). The normalized Cx43/GAPDH protein values in the human iPSC-CM samples were lower than those extracted from adult human female ventricle but substantially higher than those in negative control HeLa cells, **Figure 5G**. With natural variation of Cx43 in the samples, we saw a high degree of linear correlation to both electrical and mechanical wave CV, **Figure 5H**, confirming its critical role in conduction. The strong dependence of CV on Cx43 protein was surprising and exceeding the theoretically expected effect from pure Cx43 perturbation – 26% drop in Cx43 protein led to over 50% drop in CV for the electrical and mechanical waves. Variability in the small number of samples used and likely other parameters in the culture may have contributed to these findings, which need to be confirmed in a larger study. Nevertheless, the approach illustrates how the developed platform can be used for correlative studies to reveal molecular underpinnings of functional properties.

### Spatial patterns of the electro-mechanical delay in human iPSC-CMs

Quantifying local aEMD distribution over space can help further understand the interplay of the electrical and mechanical waves during pacing. aEMD was defined by the time lag between the onset of electrical activation (action potential depolarization) and the onset of mechanical activation (dye-free contraction). As indicated in **Figure S4**, we applied spatial segmentation based on activation patterns to boost SNR and have reliable parameter estimation, including local aEMD, along the wave path, **Figure 6A**. Segment-based action potential and dye-free mechanical traces were extracted to calculate the local aEMD. Global aEMD was simply obtained based on the space-integrated voltage and dye-free traces across the FOV. The local aEMD was shorter at the edges (at the stimulus site and the distal edge site, and higher in the middle of the samples across all pacing frequencies, **Figure 6B**. There was a strong frequency-driven increase in the aEMD both for the local (**Figure 6C**) and the global aEMD values (**Figure 6D**). Paced aEMD was longer than aEMD obtained from spontaneous waves, which also had a single origin site, **Figure 6D-E** and **Figures S7-S8**, consistent with data from global aEMD measurements in humans during sinus rhythm (shorter EMDs) vs. ventricular pacing (longer EMDs) (Pfeiffer, Tangney et al. 2014). The mechanical loading conditions in our simplified model were specific for classic cell culture – cells were firmly attached to a rigid noncompliant substrate (through extracellular matrix coating) and the edges of the dish presenting softer, more compliant attachment points due to cell growth. The shorter aEMDs at the stimulus site can be explained with better capture and synchronization, while the shorter distal aEMDs are likely due to dish-edge constraints (lower stiffness). This heterogeneity in the local aEMD is not surprising and is also seen in the wavefront itself, **Figures 3-4**. Higher frequencies exacerbate these spatial heterogeneities. These observations are consistent with previous modeling and experimental studies in whole hearts (Gurev, Constantino et al. 2010, Grondin, Costet et al. 2016). The consequences of the spatial variation of the aEMD and aEMD restitution are that conduction can be impacted in complex ways due to the interaction between the two waves; longer aEMDs can further disturb wavefront dynamics. The responses are dominated by the ECC (the mechanical wave tracks the electrical wave) but modulated by the local mechanical loading and mechanical coupling. A closer look at the local aEMD in spontaneous vs. paced activity revealed interesting dependences, **Figures S7-S8**. Samples for which the spontaneous activity site had a characteristic pacemaking-like AP, with a hyperpolarization phase (**Figure S8**), showed lower spatial APD variations and lower global aEMD, compared to sites with more ventricular-like AP with a characteristic APD “hump” (like in early after-depolarization) at the origin site. This EAD-driven wave sequences exhibited higher APD spatial dispersion (longer APD at the origin vs. distal site) and longer global aEMD, **Figure S7-S8**, consistent with clinical reports on abnormal beats, such as premature ventricular contractions, PVCs, in humans(Walia, Chang et al. 2019). Our human iPSC-CMs comprise a purified ventricular-like cell population. However, it is possible that persistent pacemaker sites may develop properties similar to sinus node drivers, which among other things, may work towards reducing the overall aEMD by establishing preferred paths of excitation. The two groups were otherwise similar in terms of their global conduction velocity.

**Figure 6.**
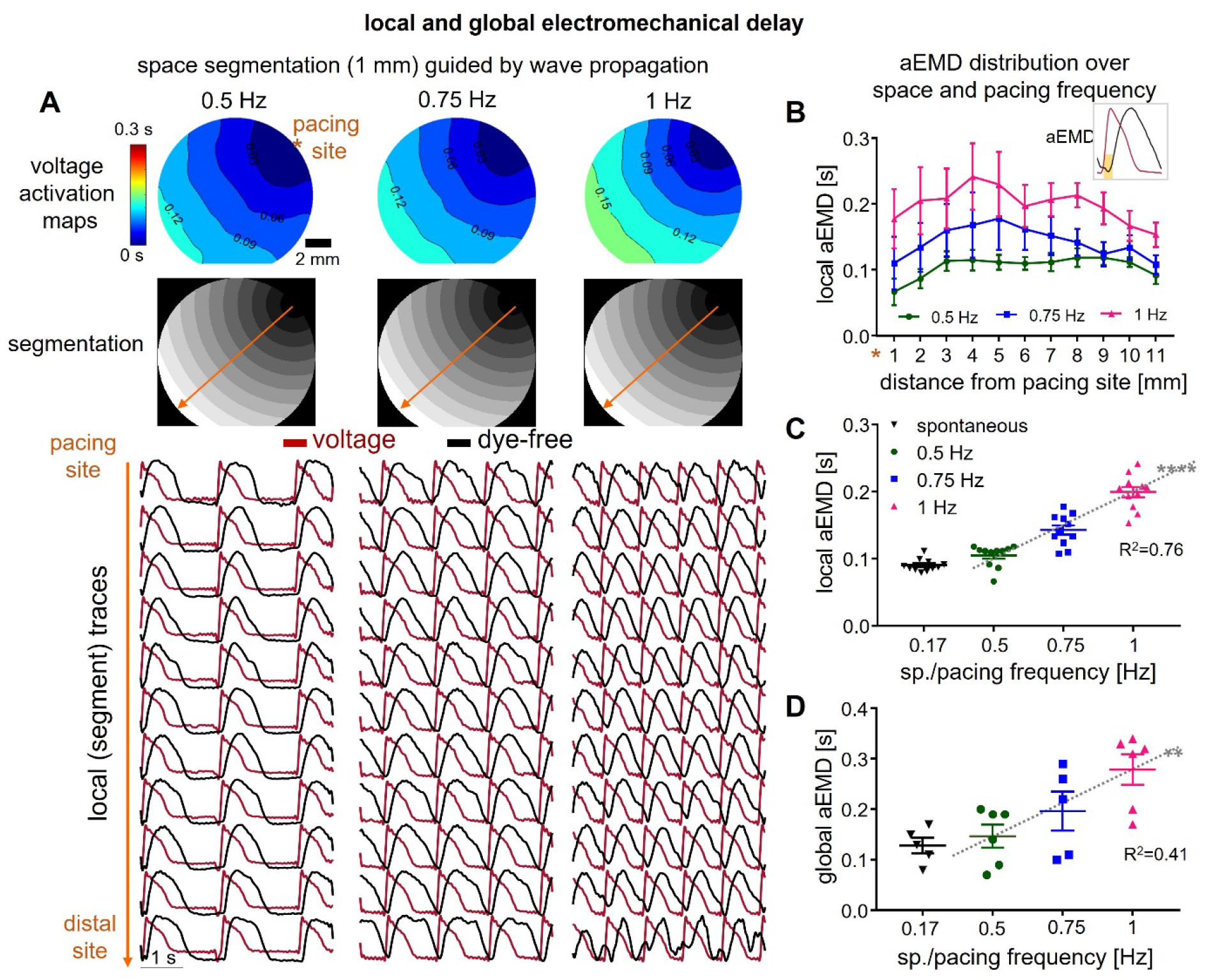
Apparent electromechanical delay (aEMD) distribution over space and pacing frequency. A. Space segmentation guided by voltage wave propagation. Shown from top to bottom: voltage activation maps, segmentation images and local voltage action potentials (red) and dye-free original traces (black) extracted from each segment under 0.5 Hz, 0.75 Hz and 1 Hz pacing. B. Local aEMD distribution over space (over the 11 segments from the pacing to the distal site) under each pacing frequency. Inset shows that aEMD was defined by the time lag between the onset of voltage action potential and the onset of dye-free contraction. C. Local aEMD as a function of pacing or spontaneous frequency (slope for pacing data: 0.19; R^2^: 0.76); shown are averaged values for the 11 segments across all samples. D. Global aEMD as a function of pacing or under spontaneous frequency (slope for pacing data: 0.26; R^2^: 0.41). Plotted data are presented with overlaid mean ± S.E.; n = 6 for pacing data and n = 5 for spontaneous data (one sample did not show spontaneous activity). The significance test for the regression lines passed at p < 0.0001 (****) for the local aEMD and at p < 0.01 (**) for the global aEMD dependence on frequency.

## DISCUSSION AND CONCLUSIONS

Parameters linking excitation and contraction in the heart are clinically important for assessing cardiac function and risk for adverse events. This study reports on the development and validation of a system for simultaneous macroscopic optical mapping of excitation and contraction waves in human iPSC-CMs, leveraging oblique transillumination, **Figure 1**. This was done with a low-cost setup, taking advantage of a method for dye-free monitoring of macroscopic contraction waves(Burton, Klimas et al. 2015), recent CMOS camera developments(Lee, Calvo et al. 2017, Heinson, Han et al. 2022), high-performance NIR voltage-sensing probes(Huang, Walker et al. 2015, Klimas, Ortiz et al. 2020) and some SNR-improving processing steps. After validating the dye-free interferometric wave mapping in our setup and showing its versatility (**Figure 2**), we characterized the properties of the observed electromechanical waves in iPSC-CM layers attached to a rigid substrate under spontaneous and pacing conditions, **Figures 3-6**, when the correspondence between two wave dynamics is relatively high. Even in this reductionist experimental model, interesting de-coupling between the excitation waves and the mechanical waves was observed as the system was “stressed” by progressively faster pacing rates. Electrical waves were more robust in their adaptation (restitution) responses, while the mechanical waves underwent major restitution-based shortening of the relaxation phase and displayed significant increase in local wavebreaks. The resulting more “tortuous” wavefronts for the mechanical waves are hallmarks of unstable pro-arrhythmic behavior, while the intact electrical waves continued to exert smoothing stabilizing action up to a point. It appeared that the areas of pronounced wavebreaks (in the contraction waves) eventually turned into areas of wavefront distortion in the excitation waves, at higher pacing frequencies, **Figures 3-4**. These interesting observations suggest utility of monitoring contraction waves as means for early detection of potential future wavebreaks and arrhythmia loci.

In general, dye-free/label-free optical mapping of cardiac waves *in vitro* is appealing as a noninvasive alternative to optical mapping with fluorescent dyes or optogenetic sensors, especially for longitudinal chronic studies using human iPSC-CMs. It is scalable, as illustrated recently for imaging of multiple samples (Ashraf, Mohanan et al. 2021). Combination with optogenetic actuation is seamless because of the flexibility with wavelength of light used, avoiding potential interferences as seen in all-optical electrophysiology(Heinson, Han et al. 2022). For maturation studies, chronic treatment, and monitoring the long-term effects of therapeutic agents, this technique can provide a surrogate view of responses, which are often closely related to the classic measures of electrical activity, as seen in **Figure 4**. Molecular correlates of cardiac wave dynamics can also be studied this way, **Figure 6**, as the dye-free/label-free wave mapping does not add new fluorescent emitters to the samples. There is a dearth of studies that offer direct links between molecular correlates and advanced cardiac responses within the same samples(Chung, Bien et al. 2011, McCain, Desplantez et al. 2012, Dhillon, Gray et al. 2013, Melby, de Lange et al. 2021), and dye-free imaging followed by gene expression (using qPCR or RNAseq) or protein quantification can fill that gap. Previously, gap junctional conductance in cardiomyocyte pairs has been shown to be linearly related to immunofluorescence-quantified gap junctional areas (when these areas are not too large)(McCain, Desplantez et al. 2012), while CV is proportional to the square root of gap junctional conductance(Shaw and Rudy 1997, Jongsma and Wilders 2000), with stronger influences in less coupled cardiac cells/tissues. No previous studies have attempted to link electrical and mechanical wave velocity to Cx43 protein content in human cardiac tissue and/or hiPSC-CMs. The significant relationships seen in this study were surprising if purely based on gap junctional coupling. Using the described pipeline, **Figure 5**, future studies will expand the sample size and conditions of plating to further dissect this relationship.

In the clinic, the state-of-the-art noninvasive technique for imaging cardiac arrhythmias is the high-density body surface electrocardiographic imaging, ECGI(Rudy 2010). However, there have been developments towards using fast ultrasound imaging to reconstruct the spatial patterns of myocardial contraction waves as a noninvasive 3D alternative of the ECGI-generated surface excitation waves. Electromechanical wave imaging (EWI) is such quantitative echocardiography-based technique(Provost, Lee et al. 2011, Grubb, Melki et al. 2020). Recent advanced electromechanical mapping using motion tracking with ultrasound imaging also provided insightful baseline comparison with optical mapping in whole hearts (Christoph, Schroder-Schetelig et al. 2017, Christoph, Chebbok et al. 2018, Christoph and Luther 2018). Leveraging machine learning techniques, computational modeling and data assimilation, efforts are underway towards the goal of reconstructing the electrical wave patterns in 3D space over time from 4D contraction wave imaging (Lebert and Christoph 2019). Such efforts promise a non-invasive clinically-relevant detailed view of cardiac arrhythmias using 4D ultrasound-based imaging. Our work on simultaneous imaging in simple systems can help validate and refine electromechanical models to be able to perform such reconstructions. Furthermore, direct testing of model-generated predictions, related to fundamental relationships between the electrical and mechanical waves, are feasible in our simplified experimental model. Because of ability to introduce controlled alterations to the substrate, such as cell anisotropy(Chung, Bien et al. 2011), compliant growth surface and a variety of deliberately created patterns of stimulation over space and time (especially with optogenetic actuation(Burton, Klimas et al. 2015)), comprehensive data sets can be generated linking the tri-wave dynamics for machine learning algorithms and improved model generation.

An example of a clinically utilized parameter linking electrical and mechanical waves is the electromechanical delay, EMD. At the cellular level, EMD reflect the ECC events, i.e. the timing from the membrane depolarization to the release of calcium from the sarcoplasmic reticulum and the engagement of the myofilaments to initiate contraction; EMD is also influenced by the mechanical loading conditions and by the specific activation sequence(Pfeiffer, Tangney et al. 2014). It has been suggested that shorter EMDs reflect stronger mechanical coupling(Nitsan, Drori et al. 2016), better synchronized contractions, and for the patient, this may mean better stroke volume and cardiac output(Weissler, Harris et al. 1968).

Longer EMDs distinguish abnormal beats – they are characteristic for EADs, a known mechanism for emergence of PVCs, compared to regular sinus rhythm beats(Walia, Chang et al. 2019). Furthermore, it has been observed that single-site ventricular pacing increases EMD heterogeneity and worsens ventricular pump performance; it may therefore present a path towards development of heart failure(Andersen, Nielsen et al. 1997, Bordachar, Garrigue et al. 2003, Kerckhoffs, Faris et al. 2005). EMD is typically prolonged in patients with heart failure(Weissler, Harris et al. 1968, Badano, Gaddi et al. 2007) and is often considered as a biomarker for seeking cardiac resynchronization therapy (CRT); lead localization for CRT can be optimized based on predicted EMDs(Constantino, Hu et al. 2013)

Clinically, EMD is assessed by differencing global parameters – the QRS timing from the ECG vs. the timing obtained from a left-ventricular pressure wave records (for example, through acoustic cardiography) or by tracking myocardial deformation (using tagged MRI). Spatial maps of EMD are nontrivial to obtain, yet valuable considering the dependence of EMD on activation sequences. Earlier low-resolution dual mapping in canines has yielded some insights about the spatial heterogeneity of EMD (Delhaas, Arts et al. 1993, Kerckhoffs, Faris et al. 2005). Recently, a combination of ECGI and cardiovascular magnetic resonance, CMR, was applied to spatially map EMD(Dawoud, Spragg et al. 2016) in patients to quantify the performance of CRT. For a finer look into spatial EMD patterns, computational modeling has been deployed (Usyk and McCulloch 2003, Gurev, Constantino et al. 2010, Gurev, Lee et al. 2011, Provost, Gurev et al. 2011, Pfeiffer, Tangney et al. 2014).

Our dual mapping approach provides new means to dissect such key parameters in a simplified, well controlled setting. For example, our culture system seems capable of distinguishing “regular” spontaneous pacing sites from sporadic, EAD-driven ones, **Figure S7-S8**, based on their different aEADs, as seen clinically(Walia, Chang et al. 2019). While we work with purified ventricular myocytes, persistent pacemaking sites may still establish preferred activation sequences, minimizing EMD when waves travel along those paths. Furthermore, faster pacing rates in our system seem to increase local and global aEMD (**Figure 6**), as seen in canine *in vivo* models, when global EMD was reported (Kettunen, Timisjarvi et al. 1985).

Yet, another parameter linking excitation and contraction, that has been useful in predicting cardiotoxicity of drugs, is the electromechanical window, EMW, defined as the difference between the duration of the mechanical event (contraction transient or left-ventricular pressure event) and the duration of the electrical trigger (APD or QT-interval). Drugs or events that lead to shorter EMWs, or to negative EMW, can be pro-arrhythmic. Clinically, EMW, based on the global ECG and left-ventricular pressure wave measurements, has been used to predict arrhythmia risk in long-QT patients(ter Bekke, Haugaa et al. 2014) and there is an ongoing clinical trial that started in 2020 (NCT04328376) based on this metric. Furthermore, experimental work in canines(van der Linde, Van Deuren et al. 2010) has demonstrated that drugs that shorten the EMW may have higher risk for arrhythmia induction, specifically risk for Torsade de Pointes. Computational work (using calcium as a surrogate for mechanical contraction) has corroborated that shortening of EMW can be useful in cardiotoxicity prediction in drug testing(Passini, Trovato et al. 2019). In our study the increase in APD/DFD ratio (**Figure 4C**) reflects a decrease in EMW in a more direct way. Therefore, simultaneous mapping of electromechanical waves, as demonstrated here, may inform cardiac safety studies with human iPSC-CMs, beyond the typically used APD prolongation as a biomarker.

In conclusion, dual mapping of electromechanical waves in human iPSC-CMs can be a useful noninvasive tool for chronic studies, can offer valuable insights about the complex dynamics between these two systems, generate comprehensive data sets for validation and improvement of cardiac electromechanical models and can offer guidance for clinical imaging, interpretation and interventions in treatment of cardiac arrhythmias.

## METHODS

### Cardiomyocyte cell culture

In these experiments, we used human iPSC-CMs-iCell2 (iCell Cardiomyocytes2 (Cat. C1016, Donor 01434) from Fujifilm/Cellular Dynamics International, Madison, WI. The cells were cultured on fibronectin-coated (50μg/mL) 35mm dishes with 14mm glass-bottom insert (Cellvis, Mountain View, CA) at 270,000 cells per dish. Samples were grown in humidified incubator at 37 □ C and 5% CO2 as per manufacturer’s instructions. In some samples, the optogenetic actuator, Channelrhodopsin2 (ChR2), was expressed using adenoviral transduction with Ad-CMV-hChR2(H134R)-eYFP (Vector Biolabs, Malvern, PA) at multiplicity of infection (MOI 50) for two days prior to experiments.

### Experimental sample preparation and protein quantification

The iPSC-CM samples were studied on day 7 after plating in Tyrode’s solution (in mM: NaCl, 135; MgCl2, 1; KCl, 5.4; CaCl2, 1.8; NaH2PO4, 0.33; glucose, 5.1; and HEPES, 5; adjusted to pH 7.4 with NaOH) at room temperature. For macroscopic mapping, samples were labeled with the near-infrared V_m_ dye BeRST1(Huang, Walker et al. 2015), courtesy of Evan W. Miller (UC Berkeley), at 1μM concentration for 20min. For some microscopic experiments, dual-labeled samples were used with spectrally-compatible fluorescent indicators for membrane voltage V_m_ and intracellular calcium [Ca^2+^]_i_ in Tyrode’s solution, as described previously(Klimas, Ortiz et al. 2020). The [Ca^2+^]_i_ indicator Rhod-4AM (AAT Bioquest, Sunnyvale, CA) was used at 10μM concentration for 20min, followed by a wash, and the application of the near-infrared V_m_ dye BeRST1 at 1μM for 20min. After a final wash, samples were imaged. Electrical stimulation was done by a bipolar platinum electrode, 2 ms pulses at 5V, connected to a MyoPacer stimulator (IonOptix, Westwood, MA). Each of the presented here samples was subjected to at least three stimulation frequencies, i.e., 0.5 Hz, 0.75 Hz and 1 Hz.

In some experiments, the uncoupling agent 2-Heptanol (Sigma Aldrich, St. Louis, MO) was applied in Tyrode’s solution at 0.5mM for 15 min prior to recordings; this concentration was chosen. to be < EC50 for cardiomyocytes (Jia, Bien et al. 2012) to avoid non-specific effects.

### Protein quantification by automated Western Blot (Wes)

After completion of the optical mapping experiments, some samples were lysed for total protein extraction using Qproteome Mammalian Protein Prep Kit (Qiagen, Hilden, Germany). Western blot was done using capillary electrophoresis based system Wes™ (ProteinSimple, Santa Clara, CA) according to the manufacturer’s instructions. The ab11370 (Abcam, Cambridge, UK) primary antibody was used in 1:25 dilution for Connexin 43 and the ab181602 (Abcam) antibody was used in 1:2000 dilution for GAPDH quantification. Detailed methods have been published(Li, Han et al. 2022). Correlative analysis was possible as the functional measures (conduction velocity) and the normalized protein (Cx43/GAPDH) were obtained in the same samples.

For the Western Blot quantification, we used as a positive control for Cx43 cell lysates, extracted from flash-frozen adult human left ventricular tissue (Amsbio, Abingdon, United Kingdom), processed as described previously(Li, Han et al. 2020). As a negative control, we used HeLa cell lysates (ProteinSimple).

### Design of a dual optical mapping system for electro-mechanical waves in iPSC-CMs

The customized simultaneous dual macroscopic imaging (SDMI) system, **Figure 1**, was built using two independent CMOS cameras (Basler_acA720-520um, Ahrensburg, Germany) with 540×720 resolution. The imaging magnification factor of the two cameras is equivalently obtained as 0.315x by an 0.15-NA objective lens (MV PLAPO 1X, Olympus, Shinjuku City, Tokyo, Japan) and a demagnification lens (LA1951, Thorlabs, Newton, MA, USA), resulting in a 1.86-cm^2^ field of view (FOV) and a 63-mm working distance. The collimated incident beam (for fluorescent excitation and scattering interference) was from a continuous monochromatic light source implemented by a 660-nm diode laser (Cobolt 06-MLD, Cobolt AB, Solna, Sweden). An aluminum-coated mirror was clockwise rotated 30 degrees to the horizontal axis for obliquely directing the light beam to hiPSC-CMs samples cultured in a dish with glass bottom. The oblique transillumination by the laser diode had a maximum irradiance of 31 mW/cm^2^ on the sample (4 samples were imaged by laser diode). An alternative light source used here was a 660-nm LED (M660L4, Thorlabs, Newton, NJ, USA) to get a higher irradiance of up to 100 mW/cm^2^ (the rest of the samples were imaged by LED). No difference in performance was registered between the two light sources. A long-pass dichroic mirror with a cut-off wavelength at 660 nm (FF660-Di02-25×36, Semrock, Rochester, NY, USA) was used for splitting the fluorescence emission light and the scattering interference light into the two cameras, respectively. An emission filter (ET595/40m+700LP, Chroma, Bellows Falls, VT, USA) was placed in the optical pathway of the fluorescence camera to suppress the excitation beam background. The lateral resolutions of the dual macroscope system were quantified using high-resolution printed letters. The fitting edge spread functions (ESF) of the sharp edges were averaged to get the line spread function (LSF). The lateral full widths at half max (FWHM) of the LSF were measured as ∼30.1 μm for both imaging modalities. Note that for dual calcium-dye free experiments, as shown in **Figure 3**, we used calcium-sensitive dye Rhod-4 at 10μ M and different light source and filter sets: a green LED (M530L4, Thorlabs) at 530nm for excitation, with excitation filter ET535/50m (Chroma), a long-pass dichroic mirror (FF552-Di02-25×36, Semrock), and the same multiband emission filter in front of camera 1 as in voltage imaging.

### Microscopic imaging setup

The microscopic imaging was done on an inverted Nikon TE 2000 microscope with a 20X objective. The cardiomyocytes were cultured in a 96-well plate with class bottom and were imaged from the bottom. The all-optical system(Klimas, Ambrosi et al. 2016, Klimas, Ortiz et al. 2020) included custom additions with 470-nm LED (M470L4, Thorlabs, Newton, NJ, USA), 530-nm LED (M530L4, Thorlabs, Newton, NJ, USA) and 660-nm LED (M660L4, Thorlabs, Newton, NJ, USA) combined into the objective for direct optogenetics pacing, calcium and voltage excitation, respectively. Another LED (M590L3, Newton, NJ, USA) with collimated yellow beam was placed on the top of the plate with an angle (45 degrees) for achieving dye-free imaging based on oblique transillumination, which avoided the excitation window of the voltage indicator. The imaging end of the system could be switched to the Photometrics 95B prime camera or the Basler camera via a rotation mirror. Optical records of voltage, calcium and dye-free contractions were collected using the two cameras under the same conditions of optogenetic pacing. While recording, the optogenetic pacing frequency was set to be 0.5 Hz with a pulse width of 10 ms, the voltage and calcium excitation irradiance was 30 mW/cm^2^, the dye-free excitation irradiance was 25 mW/cm^2^, and the frame acquisition rate of each camera was 100 Hz. Each record had a time length of 30 sec.

### Synchronized camera acquisition

The acquisitions of the voltage and dye-free cameras were implemented on a commercial USB3.0-based software Pylon Viewer (Basler). The imaging gains were 36 for the voltage camera and 10 for the dye-free camera, with a continuous excitation light irradiance of 31 mW/cm^2^ at 660 nm. The acquisition mode on the software for both cameras was set as trigger mode via detecting the external burst frame start trigger. The external synchronization TTL trigger sequences were generated by a functional generator (DG1032Z, Rigol, Beijing, China) and simultaneously delivered to the two cameras via GPIO cables. The trigger frequency was 100 Hz, resulting in 100 frames per second (fps) acquisition rate. During synchronized acquisition, the exposure time of the cameras was 9.8 ms which left 0.2 ms to allow data transmission during each triggering cycle; the recording video format was in uncompressed 8-bit AVI, and the total acquisition time were 20 sec for each pacing frequency record.

### Camera co-registration

The FOVs of the two independent cameras were calibrated by simultaneously imaging a scale card before the iPSC-CMs sample. Slightly horizontal, vertical and rotational shifts were applied for the voltage FOV to match the dye-free FOV. Three cross features scattered across the scale card images were extracted to determine the lateral shift parameters of the voltage FOV. The optimized shift parameters were globally searched until the same cross features in the two FOVs were approximately matched. The calculation process of the new registered pixel index according to those optimized shift parameters is described in supplements (for details see **Figure S3** and **Supplemental Methods**).

### Voltage and dye-free data processing

#### Temporal period-shift enhancement (PSE)

PSE takes advantage of periodic signals, and for paced records, it was applied to each pixel across all frames to enhance the signal SNR (see **Figure S4**), by iterative summation of the temporally shifted trace and the original trace, i.e.,

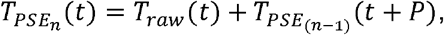

Where *T* _*raw*_ indicates the raw voltage action potential or contraction trace, 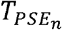 and 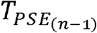 indicate the enhanced traces after *n* and *n*-1 times shift (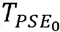 is equal to *T* _*raw*_ when *n* = 1, respectively, and *P* is the period length determined by frame acquisition rate over pacing frequency. The maximum iterative PSE number is limited to the trace length, which is given by total frame number over period. The total iterative PSE number was set to be half of the maximum in each stimulation recording.

#### Locally weighted spatial Bartlett filtering and temporal smoothing

The second SNR enhancement step was applying a same spatial-weighted average filter to all voltage and dye-free frames. We used a Bartlett filter:

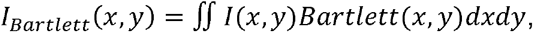

where *I* (*x, y*) indicates the original individual frame, *Bartlett* (*x, y*) indicates the 2-D kernel function which is constructed by a mesh of the 1-D Bartlett window function with a window size of 20 pixels, and *I*_*Bartlett*_ (*x, y*) indicates the spatially filtered frame. The spatial-weighted feature of the Bartlett filter prevents over-filtering and is suitable for fine wavefront tracking(Bien and Entcheva 2006, Mironov, Vetter et al. 2006). After the Bartlett filter, a temporal filter was applied by locally weighted temporal regression (“loess”) which removed the significant outliers of the traces. The span for the temporal filter was 150 ms. The purpose of temporal filtering was to clean the trace glitches for accurate detection of the event start.

#### Dye-free event start boost by time-difference (TD) filter

After PSE, co-registration was applied to the voltage FOV for precise FOV match with the dye-free FOV (see previous section). Then, a time difference (TD) filter was applied to the dye-free data for event start boosting(Burton, Klimas et al. 2015). TD filter was constructed by the absolute difference value of the PSE trace and the time-shifted trace:

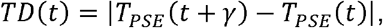

where *γ* is the difference time. It is important to mention that a TD filter may induce a phase shift in the “boosted” trace for different *γ* values, compared to the original trace due to the asymmetric shape of the transients (**Figure S4, Supplemental Methods**). This is inconsequential for constructing activation maps and quantifying conduction properties since it affects all pixels equally. However, caution should be applied when trying to extract precise local time delays linking the electrical and mechanical waves. TD filter was not used when the exact relative phase of the electrical and mechanical waves was determined, and the electromechanical delay (EMD) was calculated to avoid phase shift artifacts.

#### Activation time detection

The event start for excitation and contraction waves is marked by a rapid depolarization i.e. sharp increase in voltage or contraction force (after boosting). Based on this advantage, a time-delayed difference trace was constructed by subtracting the smoothed trace from the delayed trace, i.e.,

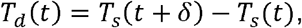

where *T*_*d*_ is the time-delayed difference trace, *T*_*S*_ is the temporally smoothed trace and *δ* is the delay. Typically, *δ* is optimized to cover the duration of depolarization, e.g. 100 ms for voltage action potential and 200 ms for contraction. The activation was obtained by recording the timing of the time-delayed difference peaks (**Figure 3A**). Activation maps were constructed as contour plots containing the event activation time of all pixels for a beat. A spatial gaussian filter with a span of 20 pixels was applied to the voltage and dye-free activation maps for further smoothing.

### CV, DFD, APD and aEMD measurements

Based on activation maps, the conduction velocity (CV) of excitation waves was calculated by the physical distance between the first ROI (near pacing/spontaneous site, see the ROI examples in **Figure 3**) and the last ROI (distal site) over the wave propagation timing in between. The chosen ROIs were matched in voltage and dye-free FOVs and kept consistent in one sample for the calculations over all pacing frequencies. Mean CV values of all samples at each pacing/spontaneous frequency were calculated and presented with S.E.

Space segmentation was first performed according to the activation maps. Each segment was approximately perpendicular to the wave propagation direction. The activation time is roughly similar within each segment, which reduced signal distortion but helps signal enhancement for spatial average. The original (without any prior spatial filtering and temporal smoothing) voltage action potential traces and dye-free transient traces within each segment were summed up and normalized. After normalization, local action potential duration, APD, was measured within each segment as the duration at 80% level (APD80, voltage action potential), local dye-free signal duration, DFD, was measured within the same segment as the time duration between the onset of contraction and the offset of relaxation (dye-free transient); local apparent electro-mechanical delay, aEMD was measured within each segment by the time lag between the onset of electrical activation (voltage action potential) and the onset of mechanical activation (dye-free contraction). We use the adjective “apparent” as the dye-free signal is assumed to be uniquely linked to mechanical contraction but may also reflect other cell properties, such as changes in index of refraction that are electrically-linked. Mean APD, DFD and aEMD values of all samples within each segment and at each pacing/spontaneous frequency were calculated.

### Wave-front tortuosity index calculation

An automated wave tracking method (in Ccoffinn(Tomek, Burton et al. 2016), based on Bayly’s approach(Bayly, KenKnight et al. 1998)) was applied to individual activation maps to generate wave-front segmentations and propagation vectors in each segment. Each segmentation frame remained activated for 200 ms. The arrow of the vectors pointed in the direction of propagation and their magnitude indicated the relative local CV. For each arrow and its neighborhood arrows within the radius of 75 pixels, the circular standard deviation (CSD) of the angular components in polar coordinates was computed. The global wave-front tortuosity index (in degrees), WT index, was then calculated by the average of the CSDs of all tracking arrows across the entire FOV.

### Statistical tests

Statistical analyses were performed using MATLAB (v.2018b) and GraphPad Prism (v.8). Data are presented as dot plots and mean +/-S.E. is overlayed. When comparing individual groups, two-tailed t tests were used to determine statistical significance. Two-way ANOVA was used for APD, DFD, CV and WT to evaluate significance among three or more groups and with appropriate post hoc tests as indicated in the text for comparisons between groups. Linear regression lines were fit and a linearity test was applied. Slopes and R squared are specified whenever possible.

## Author contributions

EE and WL conceived the study and oversaw the implementation. WL built the dual macroscope, wrote the analysis software and analyzed most of the results. JLH and WL performed all experiments. JLH prepared all experimental samples and reagents. GB advised on the dye-free imaging and camera deployment. JT analyzed wavefront tortuosity. WL and EE wrote the initial draft of the manuscript, all authors contributed to the revised version.

## Acknowledgements

We thank Evan W. Miller (UC Berkeley) for providing the NIR voltage dye Berst1 used in these studies. This work was supported by NIH grants R01HL144157, R21EB026152 and NSF EFRI1830941 to EE.

