## Supplement for "Simultaneous widefield voltage and interferometric dye-free optical mapping quantifies electromechanical waves in human iPSC-cardiomyocytes"

for

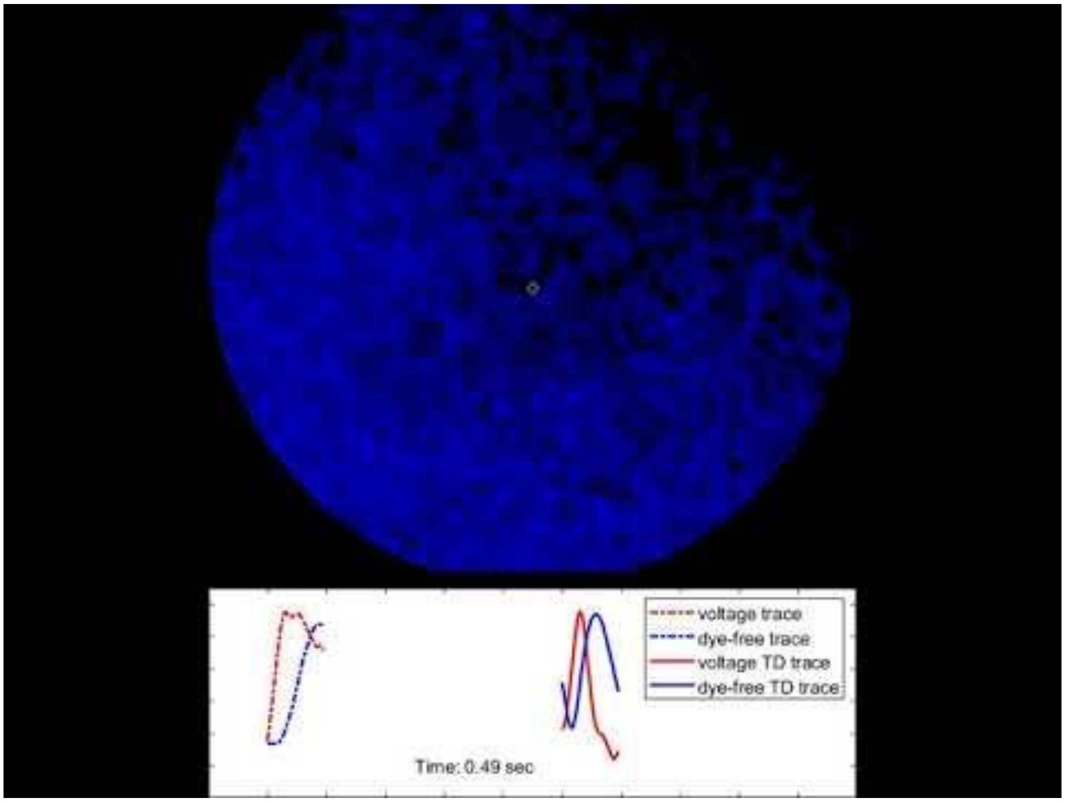


**Supplemental Video V1. Electromechanical waves under 0.75Hz pacing.** Voltage (red) and dye-free (blue) waves are shown, along with simultaneously displayed original (left) and time-difference enhanced (right) traces for voltage (red) and dye-free imaging (blue) from the indicated spot.
https://www.youtube.com/watch?v=3CIsJBRYj4M

### Supplemental Figures

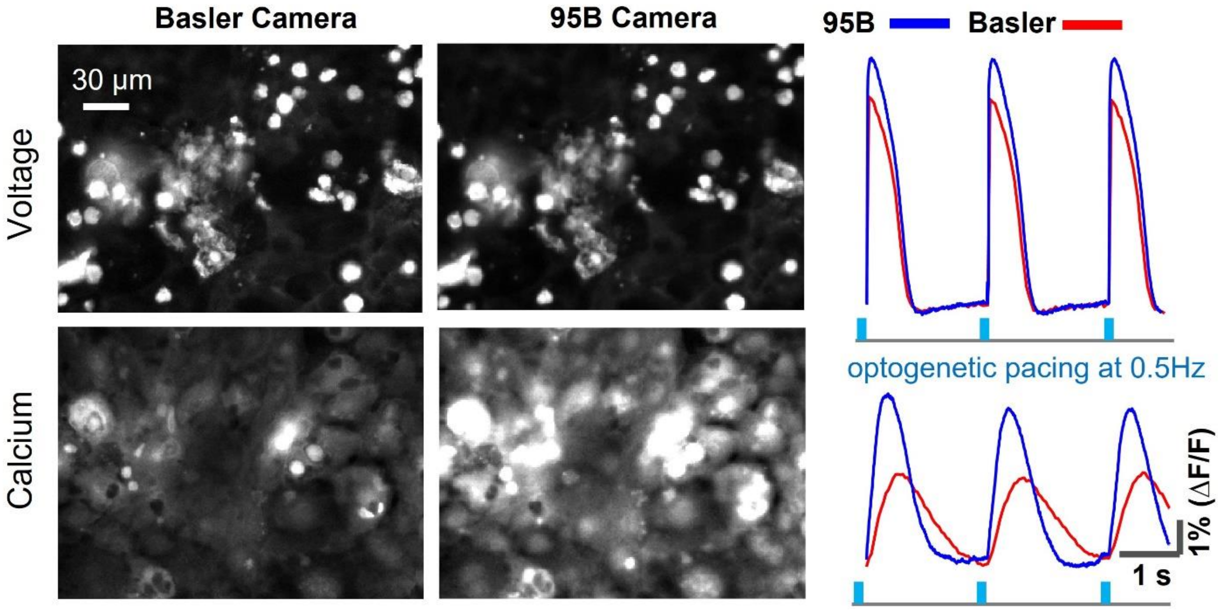


**Supplemental Figure S1**. **Simultaneous microscopic imaging of membrane voltage and intracellular calcium in hiPSC-CMs using inexpensive miniature CMOS (Basler) camera and comparison to sCMOS Photometrics 95B Prime.** Monolayers of hiPSC-CMs in 96-well format plates were infected with adenovirus to express the optogenetic actuator ChR2-eYFP and before imaging were labeled with the NIR voltage-sensitive dye BeRST1 (1μM), ex. 660nm, and the calcium-sensitive dye Rhod-4 (10 μM), ex. 535nm. The Basler ace U and the Photometrics 95B Prime were mounted on two ports of an inverted Nikon TE2000 microscope and their fields of view (FOV) were adjusted to reflect the same area. Temporally-multiplexed imaging(Klimas, Ortiz et al. 2020) of voltage and calcium onto the same camera (Basler or Photometrics 95B) was done using a 20x objective under optogenetic pacing with blue light (470nm, 10ms pulses at 0.5Hz, 0.3mW/mm$^2$); the shown signals are FOV-integrated.

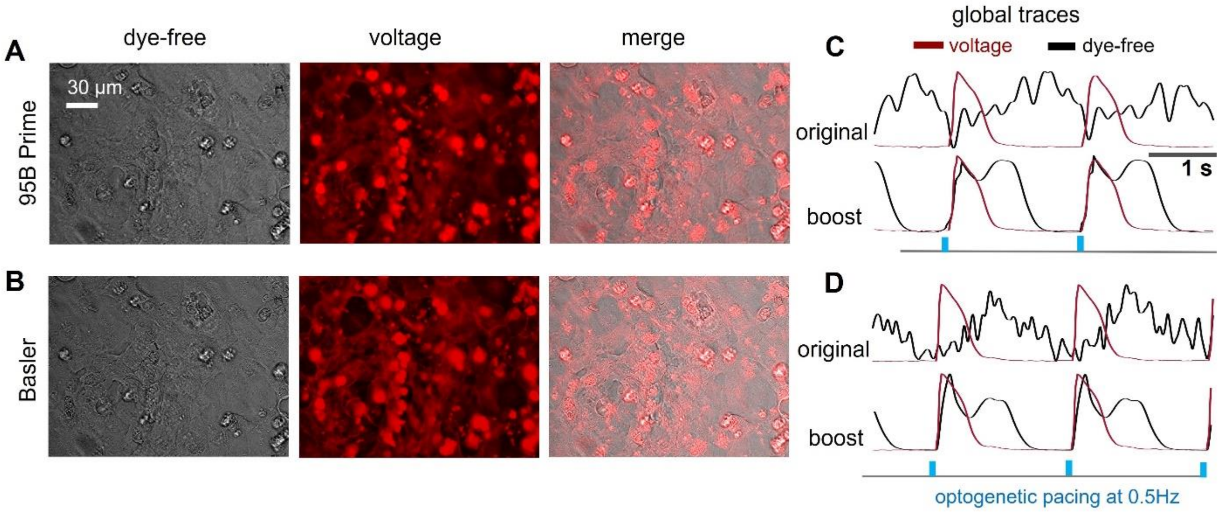


**Supplemental Figure S2**. **Simultaneous microscopic optical imaging of voltage and interferometric dye-free signal (using oblique transillumination).** Optical imaging of voltage (using BeRST1) and dye-free signal in human iPSC-CMs was done under optogenetic pacing as in **Figure S1**. Comparison of the CMOS cameras: Photometrics 95B Prime (A & C) and Basler low-cost camera (B & D). A & B. Bright-field images obtained at 590-nm excitation wavelength under oblique transillumination, voltage fluorescence images obtained at 660-nm excitation wavelength and merged images. Scale bar is 30 µm. C & D. FOV-integrated "global" traces recorded by the two cameras - voltage (red) and dye-free (black); shown are the original and the time-difference (TD)-boosted dye-free traces.

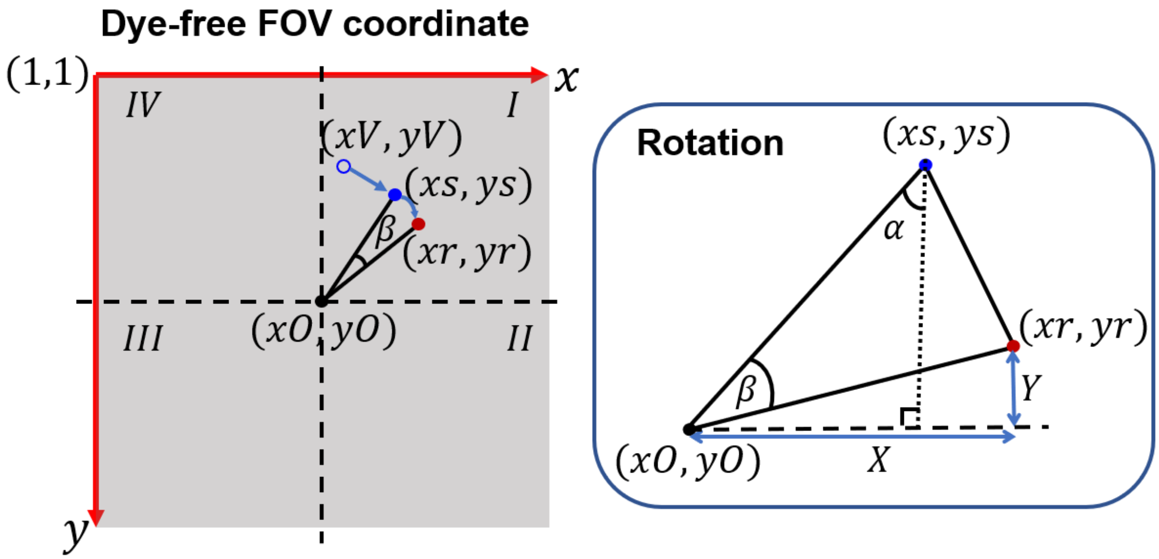


**Supplemental Figure S3. Co-registration of the voltage and dye-free cameras**. Co-registration was done by translation and rotation transforms as shown. Coordinates *xs, ys* are the pre-correction coordinates in the dye-free camera image and *xr, yr* are the corrected/co-registered co-ordinates. See Supplemental Methods for details.

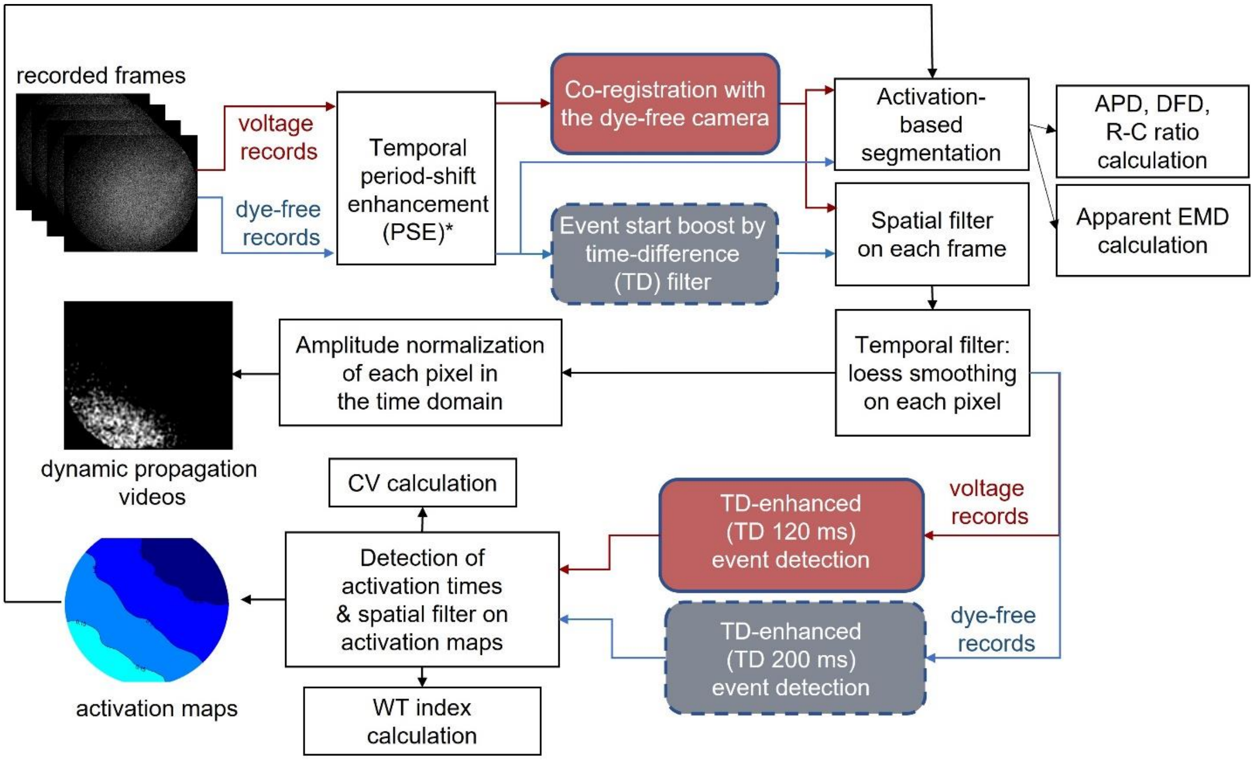


**Supplemental Figure S4. Optimized data processing for analysis of electromechanical waves from dual optical mapping of voltage and dye-free interferometric signals.** Temporal period-shift enhancement (PSE) was applied to both voltage and dye-free data at each pixel across all frames, followed by co-registration of the voltage camera with the dye-free camera and an event-start boost for the (lower SNR) dye-free signals via time-difference (TD) filter. Then, spatial Bartlett filter (span: 0.4 mm) and temporal "loess" filter (span: 150 ms) were successively applied. The dynamic wave propagation videos can be obtained at this step after normalizing the amplitude of each pixel. Activation maps were obtained for the voltage and the dye-free recordings by TD-enabled event start time detection. Spatial Gaussian filtering (span: 0.4 mm) was then applied for smoothing the activation maps. Conduction velocity (CV) and wave tortuosity index (WT) were calculated based on the activation maps. Important electromechanical parameters, such as action potential duration (APD), duration of the dye-free signal (DFD), relaxation-to-contraction (R-C) ratio and apparent electromechanical delay (aEMD) were quantified locally after space segmentation guided by the activation maps.

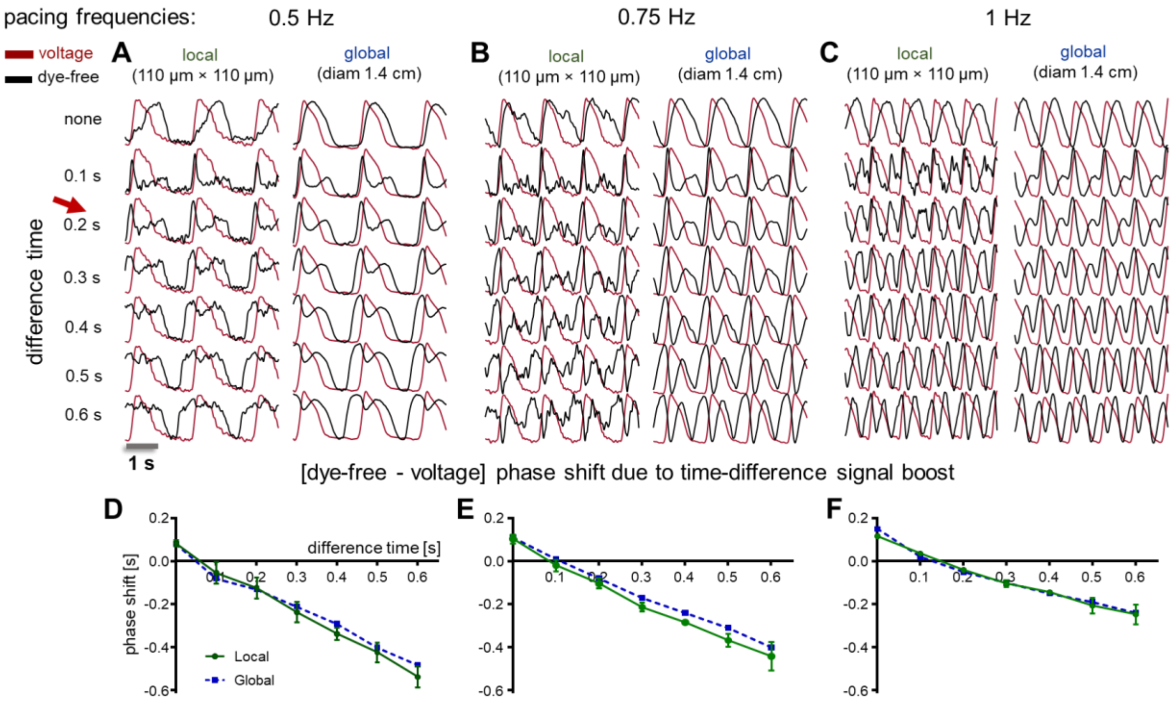


**Supplemental Figure S5. Dye-free – voltage relative position (phase-shift) analysis due to TD boost of macroscopic dye-free signals at 0.5, 0.75 and 1Hz point pacing.** A, Boosted dye-free signals (black), obtained by time difference (TD) shift from no shift (original trace) to 0.6s shift, plotted along with region-matched voltage traces (red) stimulated at 0.5 Hz. Shown are the “global” (integrated over the 1.4cm FOV) signals and “local” (110 x 110 um area) signals. B-C. Same as in A - boosted dye-free traces at 0.75 Hz and 1 Hz pacing. D-F. Dye-free – voltage phase shift (s) due to time-difference signal boost – shown are local (green) and global (blue) shifts; red arrows indicate the typically used 0.2 s TD. Data are presented as mean±S.E for n = 3 ROIs (region of interest) from a representative sample.

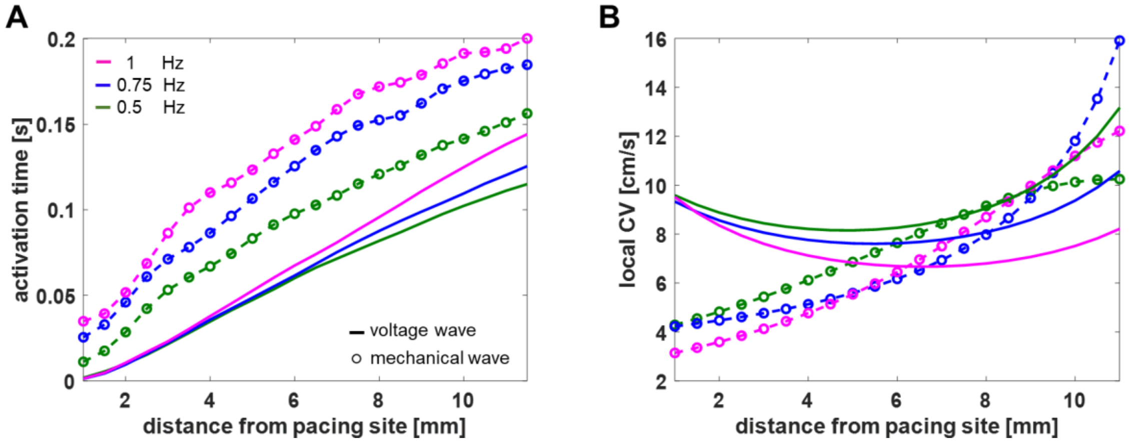


**Supplemental Figure S6. Activation time and local conduction velocity distribution along the excitation wave propagation direction in a chosen sample based on segmentation analysis.** The local aEMD analysis in Figure 6 has indicated that aEMD was not uniformly distributed along the excitation wave propagation direction. This can be due to the inconsistent local conduction velocities of the voltage and mechanical waves during propagation or due to variable mechanical loading conditions. For simple validation, activation time distribution was firstly calculated based on the same segmentation analysis method (Figure 6). Panel A shows the activation time as a function of wave propagation distance. After polynomial fitting of the activation time versus propagation distance using a least-squares algorithm, the local conduction velocity was estimated by the gradient of activation time. B shows the local conduction velocity distribution, which clearly indicates a speed up for the mechanical wave from the pacing site to the opposite edge of the dish; it becomes faster than the voltage wave before arriving at the distal end. The consequence of this type of conduction velocity distribution is the spatial variation of the aEMD – shorter at the stimulus site and at the distal border but higher in the middle, as shown in Figure 6B.

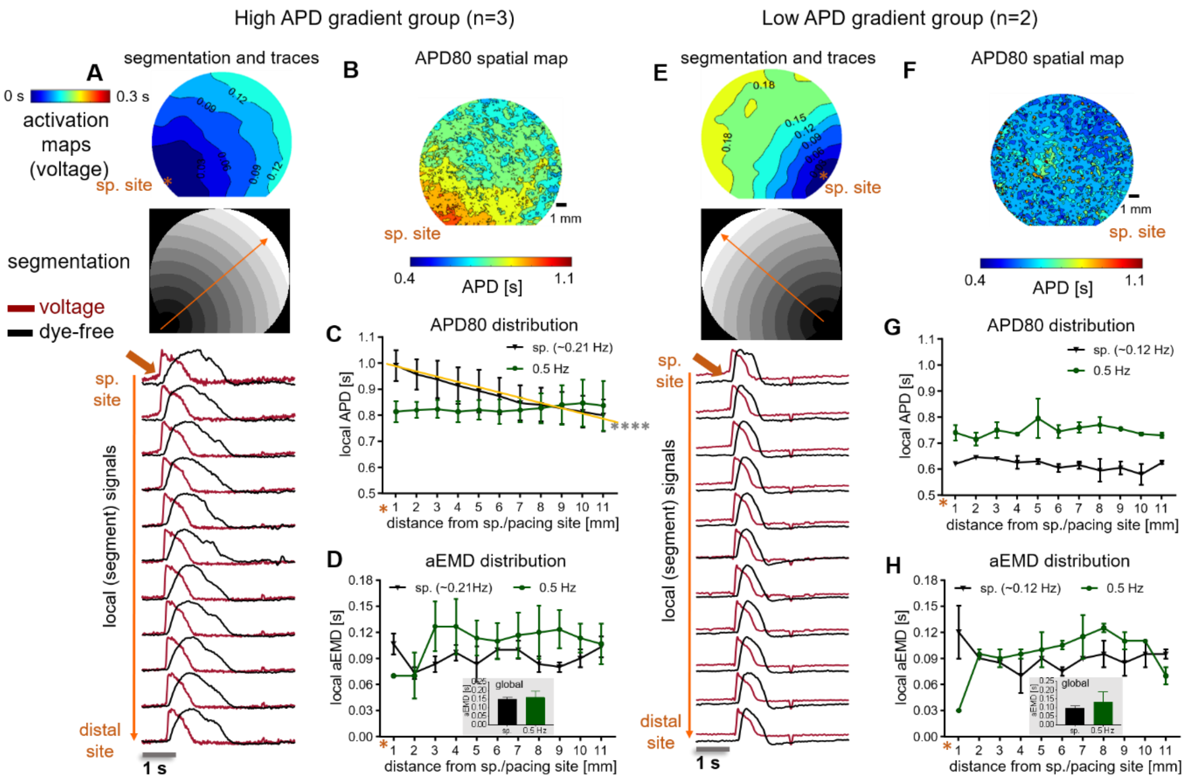


**Supplemental Figure S7. Spatial analysis of APD and aEMD for hiPSC-CMs with different APD gradients during spontaneous (non-paced) activity.** In analyzing apparent electromechanical delay (aEMD), we noted that there were two groups in terms of spontaneous activity – one with high-APD gradient (n=3) and one with low-APD-gradient (n=2); one of the 6 samples analyzed did not have spontaneous activity at the time of the measurements. A. Space segmentation guided by voltage wave propagation for high APD gradient group. Shown from top to bottom: voltage activation maps, segmentation image and local voltage action potentials (red) and dye-free original traces (black) extracted from each segment. B. Spatial APD heterogeneity MAP of one sample with high APD gradient. C. Comparison of space distribution of local APD for high APD gradient group during spontaneous rhythm and pacing (spontaneous APD gradient slope: -0.06; $R^2$: 0.21). D. Comparison of space distribution of local aEMD for high APD gradient group during sinus rhythm and pacing. Inset shows global aEMD during spontaneous rhythm and 0.5-Hz pacing. E-H, the same analysis for low APD gradient group. Data are presented with overlaid mean ± S.E.; significant linear trend is indicated by the gold line with a significance flag ($p<0.05$); n = 5.

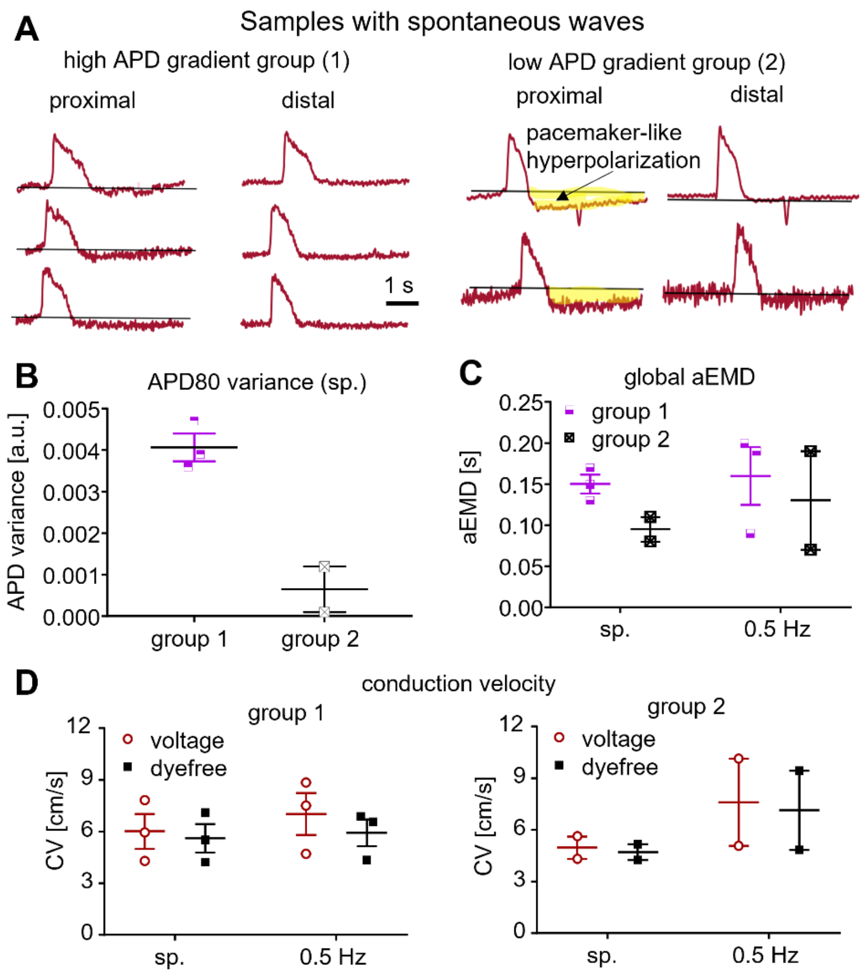


**Supplemental Figure S8. Analysis of electromechanical parameters for samples with different APD gradients during spontaneous rhythm.** Further analysis for the two groups presented in **Figure S5**. A. Example voltage traces from the proximal and the distal site for the two groups. Note the longer APDs at the site of spontaneous activity and the APD shortening at the distal site for group 1. Note the pacemaker-like hyperpolarization at the pacing site but lack of such hyperpolarization distally for group 2. B. APD80 for group 1 is longer than APD80 for group 2. C. The global aEMD for group 1 is higher than that for group 2 for spontaneous rhythm and for 0.5Hz pacing, despite similar conduction velocities during spontaneous activity and during 0.5 Hz pacing in the two groups, shown in panel D. Shorter global aEMDs (as in group 2 with the hyperpolarization-displaying pacemaking site) may reflect better mechanical coupling(Nitsan, Drori et al. 2016); while the higher APD-gradient group, where spontaneous activity may be a result of early or delayed after-depolarizations shows longer global aEMDs and therefore potentially lower mechanical coupling.

### Supplemental Methods

#### Co-registration of voltage and dye-free imaging cameras in widefield optical mapping

Please see **Suppl. Figure S2.**

Assuming $(x_V, y_V)$ indicates the pixel index in the voltage frame, after lateral shift, the index of the voltage frame registered in the dye-free camera coordinate becomes $(x_S, y_S)$ which is given by:

$$x_s = x_V + HS,$$

$$y_S = y_V + VS,$$

where $HS$ is the horizontal shift and $VS$ is the vertical shift. After further rotation around the dye-free FOV center $(x_O, y_O)$, the registered pixel index is indicated by $(x_r, y_r)$, as shown in **Figure S2**; the close-up shows the calculation principle of $(x_r, y_r)$. Here, the angle α is constructed by the connection line between $(x_S, y_S)$ and $(x_O, y_O)$ and the vertical line from $(x_S, y_S)$ to the most closest axis in the counterclockwise direction, which can be obtained as

$$\alpha = \begin{cases} \arctan(|x_S - x_O|/|y_S - y_O|), \ if \ (x_S, y_S) \in I, III \\ \arctan(|y_S - y_O|/|x_S - x_O|), \ if \ (x_S, y_S) \in II, IV \end{cases},$$

where $I, II, III$ and $IV$ represent the different coordinate quadrants. As the distance between $(x_O, y_O)$ and $(x_S, y_S)$ is equal to that between $(x_O, y_O)$ and $(x_r, y_r)$, so the horizontal and vertical displacements of the registered pixel to the center pixel are given as:

$$X = \begin{cases} \sqrt{((x_S - x_O)^2 + (y_S - y_O)^2)} sin(\alpha + \beta), \ if \ (x_S, y_S) \in I, III \\ \sqrt{((x_S - x_O)^2 + (y_S - y_O)^2)} cos(\alpha + \beta), \ if \ (x_S, y_S) \in II, IV \end{cases},$$

$$Y = \begin{cases} \sqrt{((x_S - x_O)^2 + (y_S - y_O)^2)} cos(\alpha + \beta), \ if \ (x_S, y_S) \in I, III \\ \sqrt{((x_S - x_O)^2 + (y_S - y_O)^2)} sin(\alpha + \beta), \ if \ (x_S, y_S) \in II, IV \end{cases}.$$

Finally, the registered pixels are expressed as:

$$x_r = \begin{cases} x_O + X, \quad if \ x_S \in I, II \\ x_O - X, \quad if \ x_S \in III, IV \end{cases},$$

$$y_r = \begin{cases} y_O + Y, \quad if \ y_S \in II, IIV \\ y_O - Y, \quad if \ y_S \in I, IV \end{cases}.$$

**Phase shift analysis of event start-boosted dye-free signals by time-differencing (TD)**

Please refer to **Figure S5**.

For symmetric sinusoidal signals, the time-differencing method for better event detection ("event-start-boosting") works without introducing distortions. However, for complex-shape asymmetric signals, like the voltage or dye-free signals here, certain phase shift can be introduced by this approach.

We analyzed this phase shift for data over all pacing frequencies and for TD shift from zero to 600 ms. Local phase shifts were measured in at least three different regions (110 μm × 110 μm) to obtain an average and global phase shifts were measured from the entire FOV. Statistical analysis for the local phase shift was performed. The dye-free event start in each trace can be boosted after applying TD, which is essential for determining the activation time for low-SNR data sets.

To avoid phase shift artifacts, when quantifying the apparent electromechanical delay (aEMD) in this work, we used signals without TD boost, as per **Figure S4**.
